# Radiation Exposure Determination in a Secure, Cloud-Based Online Environment

**DOI:** 10.1101/2021.12.09.471993

**Authors:** Ben C. Shirley, Eliseos J. Mucaki, Joan H.M. Knoll, Peter K. Rogan

**Affiliations:** CytoGnomix Inc., 60 N. Centre Rd., London Ontario N5X 3X5, Canada; Dept. Biochemistry, University of Western Ontario, London Ontario N6A 3K7, Canada; Dept. Pathology and Laboratory Medicine, University of Western Ontario, London Ontario N6A 3K7, Canada

## Abstract

Rapid sample processing and interpretation of estimated exposures will be critical for triaging exposed individuals after a major radiation incident. The dicentric chromosome (DC) assay assesses absorbed radiation using metaphase cells from blood. The Automated Dicentric Chromosome Identifier and Dose Estimator System (ADCI) identifies DCs and determines radiation doses. This study aimed to broaden accessibility and speed of this system, while protecting data and software integrity. ADCI Online is a secure web-streaming platform accessible worldwide from local servers. Cloud-based systems containing data and software are separated until they are linked for radiation exposure estimation. Dose estimates are identical to ADCI on dedicated computer hardware. Image processing and selection, calibration curve generation, and dose estimation of 9 test samples completed in <2 days. ADCI Online has the capacity to alleviate analytic bottlenecks in intermediate-to-large radiation incidents. Multiple cloned software instances configured on different cloud environments accelerated dose estimation to within clinically relevant time frames.

## INTRODUCTION

Despite advances in computer-assisted dicentric chromosome (DC) recognition in metaphase cell images^(1)^ and workload sharing^(2)^, quantification of radiation exposures remains a labour-intensive bottleneck in radiation biodosimetry. With few exceptions^(3)^ the same microscope system performs both data capture as well as image analysis, during which time it is unavailable for additional samples. After a large-scale radiation incident, these issues may delay timely reporting of significant overexposures in some individuals who might require treatment. Semiautomated analytic approaches requiring manual review^(4)^ will also delay reporting timely results. Outsourcing DC analyses to the fully Automated Dicentric Chromosome Identifier and Dose Estimator System (ADCI) on a dedicated high-performance computer could significantly increase overall throughput^(1,5)^. ADCI selects suitable metaphase images^(6,7)^, detects DCs^(8,9)^, generates calibration curves and estimates whole or partial-body radiation dose in samples with unknown exposures^(5,10)^. Other software programs^(11)^ can estimate exposures but do not detect Giemsa-stained DCs.

Population-scale exposures with datasets derived from *ex vivo* irradiated samples on a supercomputer^(1)^ substantially increase ADCI’s image processing speed, however these systems are not widely available to radiation biodosimetry laboratories. To achieve similar or faster results, we analyze samples with an array of lower throughput, globally available, cloud computer systems. Cloud computing outsources economically accessible computing hardware, storage, or software on-demand. These resources can accommodate both intermittent and burst throughput requirements. Similar computing architectures have been implemented in other healthcare-related domains, including deep learning, bioinformatic, and digital pathology applications^(12,13)^. Cloud-based resources would be useful in radiation cytogenetic biodosimetry because they can be allocated based on the workload, e.g., the number of samples and metaphase cell images analyzed.

The advantages of cloud computing must be balanced against cybersecurity risks. Data security and regulatory considerations must be addressed during system design^(14)^. Implementation of safeguards can avoid breaches from theft, human error, hacking, ransomware and misuse^(15)^.

In this paper^(16)^, we implement ADCI within a secure zero-trust, cloud computing framework available from Amazon Web Services (AWS)^(17)^. ADCI Online can be accessed through AWS AppStream 2.0, a fully managed streaming service for Desktop software. Separation and encryption of sensitive data are key to cloud-based data protection^(18)^. Users upload locally stored metaphase cell images to an AWS Simple Storage Service (S3) bucket which is subsequently linked to ADCI within an interactive streaming session. ADCI Online remotely enables rapid estimation of radiation exposures of uploaded samples, extending the capacity of a physical workstation with ADCI installed.

## METHODS

MS-Windows^®^-based ADCI has been ported to AWS AppStream 2.0, which is accessible worldwide through local AWS nodes. Operationally, ADCI Online is indistinguishable from the version that runs on a dedicated, standalone computer. The cloud-based system was configured by default with lower throughput hardware (2 x Intel(R) Xeon(R) CPU E5-2686 v4, 3.75GB RAM), which was ~3 fold slower than the high performance computer on which ADCI was previously benchmarked (Intel i7-6700HQ, 16GB RAM)^(1,5)^.

### On-demand remote access

The cloud version of ADCI software runs on different streaming instances, each instance having independent capabilities to process full sets of biodosimetry data. During each ADCI Online streaming session, one or more streaming instances are cloned from a master base software image or “snapshot” that was created with the AWS Image Builder tool. The snapshot is comprised of a MS-Windows^®^ Server 2016 system with ADCI preinstalled. ADCI did not need to be recompiled for this purpose and the standard installation package used for MS-Windows desktop computers was used to install ADCI on the cloud-based Image Builder tool. Snapshots can be used to clone streaming instances with distinct computer hardware configurations belonging to either the “General Purpose”, “Compute Optimized”, and “Graphics G4dn” instance families (described in https://doi.org/10.5281/zenodo.5761745). These included “stream.standard.medium” (Standard) and the faster (G4) Graphics Processing Unit “stream.graphics.g4dn.xlarge” template (Graphics G4dn) AppStream 2.0 configurations. The same ADCI snapshot was used to clone streaming instances from the “General Purpose” and “Compute Optimized” families. A second snapshot was built on the G4 template to clone Graphics G4dn instances. By default, a streaming instance boots using the Standard hardware configuration. Although less powerful than a high-performance MS-Windows^®^ system running ADCI, the cloud-based design of ADCI Online allows for rapid expansion of computing resources (Fig. 1).

**Fig. 1.**
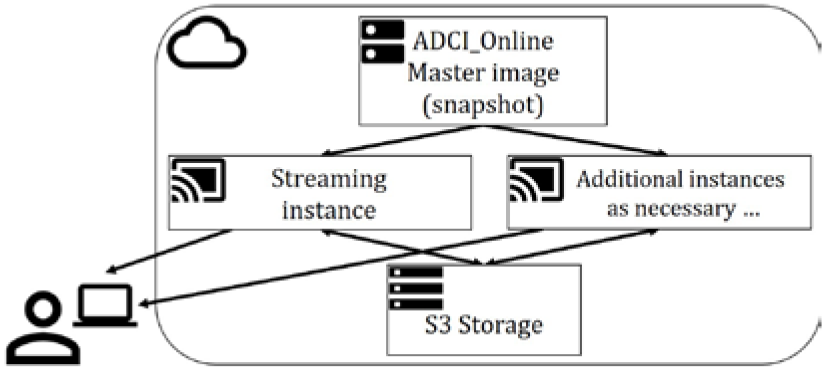
ADCI Online leverages a cloud-based environment to expand system resources on-demand. All streaming instances are configured identically from an existing snapshot and share the same permanent S3 bucket that contains metaphase images, ADCI outputs and reports. Throughput can be expanded by: (1) cloning system snapshots to provide an array of streaming instances, and/or (2) expanding these computing resources by leasing higher capacity hardware.

### Interacting with ADCI Online

Before a streaming session begins, a user interfaces with their S3 folder using the Data Storage and Retrieval web app based on the AWS Javascript Software Development Kit (SDK) v2.977.0 (https://doi.org/10.5281/zenodo.5761745). The web app is accessed with a web browser (HTTPS protocol) and implements client-side Javascript that uploads user metaphase cell images to their S3 folder via supplied AWS Cognito credentials (Fig. 2). Uploads are verified by displaying image counts of each sample. Users can view uploaded content, delete metaphase images, and download ADCI reports after they are generated. AWS currently limits the connection between AppStream 2.0 instances and S3, allowing only the first 5,000 files in an S3 directory to be accessed by the streaming instance. In some cases, this limit might impact results from ADCI Online, for those samples consisting of > 5,000 cell images. During uploading of cell images, the Data Storage and Retrieval web app recognizes samples surpassing this file limit and overcomes it by automatically directing file uploads to linked subfolders created within the new sample directory on S3.

**Fig. 2.**
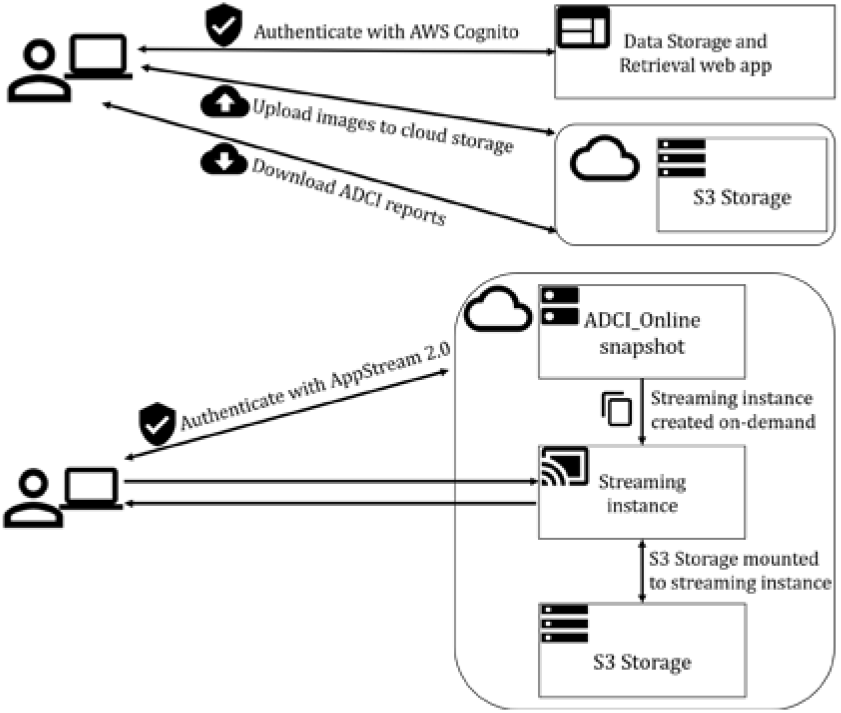
User interactions with ADCI Online. Top: Once a user validates their AWS Cognito credentials, they sign into the Data Storage and Retrieval web app to upload metaphase images to cloud storage (S3). Bottom: Afterwards, they sign into AWS AppStream 2.0 to request a new ADCI Online streaming session. The user specific S3 folder is mounted to the streaming instance, allowing the user to access their uploaded metaphase images, process samples and save results. Top: After sample processing, dose estimation and report generation, users access and download reports with the Data Storage and Retrieval web app.

Upon image transfer, credentials to access streaming instances are provided through another web link. During a streaming session, one or more streaming instances are cloned from the master snapshot of the ADCI Online system. Simultaneously, the user’s persistent S3 folder is mounted to each cloned instance, Streaming instances are controlled independently by the user, and each instance presents a copy of ADCI enabled to access a shared pool of metaphase images, ADCI outputs and reports stored in the user’s persistent S3 folder. Each instance is functionally equivalent to ADCI on the user’s local computer, including the ability to manually curate images, if desired.

The user may log out of a session, which retains all processed image files and results should a new session be activated. AWS requires that the maximum duration of the session be specified. To avoid prematurely terminating image processing, both the length of each streaming session and the disconnect timeout window have been extended to the maximum duration permitted (96 hr). Loss of internet connectivity will not discontinue processing samples within this timeout window. However, a new streaming session will reinitialize the session timer. During sample processing, files specifying chromosome contours are generated for each image. These files, which indicate chromosome boundaries and classification, are initially placed in the streaming instance’s temporary storage, then later archived, compressed, and copied to the S3 bucket upon saving a processed sample. Upon loading a processed sample from the S3 bucket, the same files are extracted and placed in the streaming instance.

### DC assay samples analyzed with ADCI Online

Typical laboratory intercomparison datasets consisting of calibration (10 samples, 50,497 images) and test (6 samples, 4,098 images) samples obtained from Health Canada (HC)^**(5)**^ and Public Health England (PHE)^**(10)**^ were uploaded to and processed by ADCI Online. Calibration curves were generated, and wholebody exposures of HC samples were determined^(5)^. Partial-body exposure levels were estimated for PHE samples, with each receiving the equivalent of 50% fractional exposures, based on an associated dose calibration curve^**(10)**^.

### Overview of ADCI functionality

Uploading of metaphase images and determination of radiation dose are illustrated online in a Supplementary Document (https://doi.org/10.6084/m9.figshare.17129768). ADCI follows IAEA guidelines for radiation biodosimetry^(19)^. ADCI processes calibration and test samples of metaphase cell images originating from the same laboratory and identifies DCs in both sample types^(8,20)^. Image selection models consist of digital filters and image ranking protocols to include images with optimal cell and chromosome morphologies^(6)^. Model optimization searches calibration samples for image selection model parameters, ranking generated models by p-value of Poisson fit, curve fit residuals, or leave-one-out cross validation^(5)^. DC frequencies in processed calibration samples filtered with an image selection model were used to fit a calibration curve using the maximum likelihood method^(21)^. The same image selection model is applied to processed test samples of unknown exposures^(9)^. Whole-body dose is estimated from the DC frequencies of test samples using a linear quadratic equation fit to the calibration curve. Partial-body dose and fraction of blood irradiated is determined with the Contaminated Poisson method after adjusting for false positives identified in unirradiated controls^(10)^. ADCI generates user reports describing samples, calibration curves, image model optimization and dose estimation, which can be downloaded from S3 storage. Details are available in the ADCI manual (https://adciwiki.cytognomix.com).

### Security and data management

An S3 bucket permanently stores uploaded metaphase images and ADCI results, and persists between streaming sessions. Each user is assigned to a unique folder in S3. Access to folders created for other users is disabled with user-specific AWS Identity and Access Management (IAM) Policies. By default, all access to the S3 bucket is disabled and each permissible action must be explicitly granted by the IAM policy, therefore if an IAM policy fails to attach to a user for any reason, they will not be granted access. All persistent ADCI Online user files are encrypted in transit to and from S3 storage (HTTPS protocol) and server-side encryption is applied to all files in the bucket. Although sensitive information may exist in an unencrypted state during streaming while actively in use, AppStream 2.0 instances exist only for the duration of a password-protected streaming session. New streaming instances execute ADCI in a clean environment that cannot be influenced by other users (preventing sensitive data or potentially malicious files from being carried forward). When the session is completed, the streaming instance and its associated temporary file storage (Amazon Elastic Block Store) is deleted.

The streaming instance is not internet accessible to third parties, cannot interact with the user’s local file system or external devices, and files can only be transferred to/from the streaming instance by way of the S3 bucket. The user-specific folder in the S3 bucket is mounted to the cloud-based system while the streaming session is active. A Microsoft AppLocker policy disallows execution of all software or scripts except “whitelisted” software directly related to ADCI or AppStream 2.0. Real time outputs are encrypted, and pixels streamed directly to the user. ADCI Online does not consume computing resources or store data on the user’s local computer system. This prevents the inadvertent transfer of sensitive data to the local system, keeping it self-contained in S3. It is also a valuable layer of application-level security, as user interaction with ADCI is limited to software-curated keyboard and mouse inputs only.

Uploaded metaphase image data are created by, accessible to and removable by individual users. Programs automatically delete user data one month after uploading and processing, unless long term archiving is requested by the user. Uploaded files and headers of uploaded images are verified as either TIFF or JPG format. Sample or report files with prohibited characters in MS-Windows^®^ file names are also disallowed.

## RESULTS

### Cloud software validation

Image processing, image selection model optimization, calibration curve generation, and dose estimation capabilities of ADCI Online were compared to output generated on a dedicated MS-Windows^®^-based computer system and to previously published studies of the same samples. HC samples were uploaded and processed on the Standard and Graphics G4dn hardware configurations. Dicentric chromosome counts and frequencies for these samples were identical with both ADCI Online and the MS-Windows^®^ version of ADCI running on local computer hardware. The best performing image selection models were identical (model scores, image exclusion filters, and image ranking methods).

Radiation exposures of HC and PHE samples were estimated with ADCI Online. Dose estimates for whole-body HC test samples were generated for 4 image selection models, matching previous results^(5)^ (Table 1). Estimates of partial-body dose and fraction of blood irradiated by PHE were largely concurrent with prior findings^(10)^. Small variations in the results of partial-body analysis in ADCI are expected due to iterated random image sampling that is carried out to correct for false positive DC^(10)^. This resulted in small differences (< 0.1%) in the estimated fractions of blood irradiated for PHE samples E, F, and G.

**Table 1.** Estimated exposures with ADCI Online.

| Sample | Physical |  | Estimated |  |  |
| --- | --- | --- | --- | --- | --- |
|  | Dose (Gy) | Frac.* (%) | Total body dose | Partial body dose | Frac. |
| HC01<sup>1</sup> | 3.1 | 100 | 3.3<br>[1.9, 6.0]<sup>3</sup> | - | - |
| HC04 | 1.8 | 100 | 1.8<br>[0.7, 3.4] | - | - |
| HC05 | 2.8 | 100 | 3.3<br>[1.9, 6.0] | - | - |
| HC07 | 3.4 | 100 | 2.9<br>[1.6, 6.0] | - | - |
| HC08 | 2.3 | 100 | 2.5<br>[1.3, 6.0] | - | - |
| HC10 | 1.4 | 100 | 1.0<br>[0.0, 2.4] | - | - |
| PHE_E<sup>2</sup> | 4.0 | 50 | 1.6<br>[0.9, 2.4] | 5.2<br>[4.0, 6.2] | 47.7<br>[31.5, 67.2] |
| PHE_F | 2.0 | 50 | 1.4<br>[0.6, 2.2] | 2.2<br>[0.9, 3.2] | 65.8<br>[35.7, 100.0] |
| PHE_G | 6.0 | 50 | 1.5<br>[0.8, 2.3] | 3.5<br>[2.3, 4.5] | 52.4<br>[31.8, 81.0] |

\*Frac. = fraction irradiated; <sup>1</sup>Sample identifiers and image selection model A\_B are described in reference 5; <sup>2</sup>Sample identifiers and image selection model C\_B750 are described in reference 10; <sup>3</sup>Bracketed values indicate 95% confidence intervals.

### Uploading and image processing performance

A set of 16 HC samples containing 54,595 metaphase images (41.7GB) was uploaded to cloud storage in 53.6min through our web app implemented using the AWS Javascript SDK. The Data Storage and Retrieval web app uploads files concurrently in batches. The HC 0.25Gy calibration sample (11,896 images) was uploaded in 11.1min in batches of 20 images. The same sample was uploaded for batches of 30 in 7.5min. Faster network connections with high-performance computers can accelerate uploads by the user by increasing the batch size parameter. All uploaded HC samples were processed on Standard and on Graphics G4dn instances. All samples completed processing with a single Standard instance in 46.5hr and in 14.4hr with a Graphics G4dn instance. For comparison, samples were split into 5 subsets with similar image counts (10,520 to 11,896 images) and processed in parallel by 5 Standard instances (Standard x5). Processing of images from individual samples is not distributed among multiple instances, in contrast with the supercomputer implementation of ADCI^(1)^. Processing with the parallelized Standard instances ranged from 7.8 to 9.3hr in length.

### Speed of overall process

Process times for calibration and test samples, optimizing image selection models, exposure determination and other operations are indicated in Table 2. Calibration sample processing and image selection optimization steps are performed only once for each laboratory, and are used for analysis of whole or partial body exposures of test samples during the same (or reused in a future) session. Images were processed at rates of 19.58 and 97.53 images/min, respectively, for Standard and Standard x5 instances.

**Table 2.**
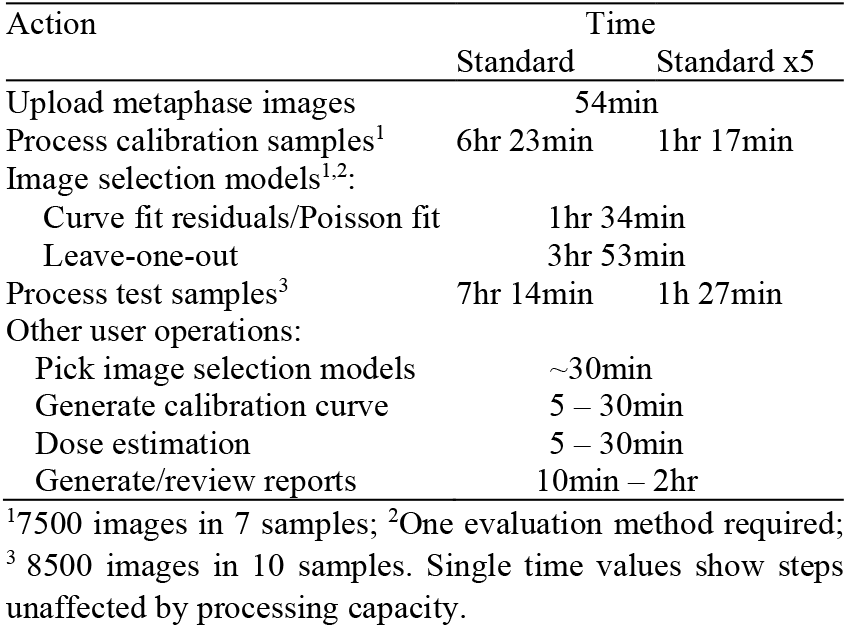
Session times for Standard hardware instances.

The time required to process samples and generate optimal image selection models is based on hardware performance. Time estimates can be obtained from processing rates (images/min) for specific hardware configurations and the number of metaphase images. “Other user operations” require little to no processing time. The duration of these steps varies according to the operator’s knowledge of the system and the volume of test samples to be analyzed. The time required to perform all steps on a single Standard instance was 17hr 11min (or 19hr 30min for leave-one-out image selection model evaluation), including 2hr for “Other user operations”. The time required for Standard x5 instances to process these samples was reduced to 6hr 32min, or 8hr 51min by leave-one-out evaluation. Based on these results, it should be feasible to complete the interpretation of typical intercomparison biodosimetry datasets within 1-2 days.

## DISCUSSION

Radiation dose estimation with ADCI Online requires only metaphase cell images from potentially exposed individuals and a reliable internet connection. We previously showed that analysis of 1,000 samples on ADCI in a supercomputer environment was ~10 fold faster than semi-automated review and analysis^(1)^. ADCI Online can also produce timely dose estimates in an intermediate to large-scale nuclear event. Should local internet access become unreliable during a radiation emergency, sample images could be exported from automated microscope systems to portable storage and either uploaded to AWS via satellite internet or transported to locations with reliable connectivity.

Image processing can be accelerated either by opting for more expensive, high performance GPU hardware or by cloning more instances, thereby increasing the level of parallelization. Multiple instances of Standard and/or Graphics G4dn instances are as efficient and cost-effective as more powerful and costly memory configurations, which process images and generate result matrices, but do not increase sample throughput. In general, it is more cost effective to clone additional copies of Standard instances rather than opting for more expensive hardware. We have demonstrated that G4 instances process metaphase images ~3.2 times more quickly than Standard instances, however the hourly AWS charge for G4 instances is currently 10 times higher. In a large-scale radiation incident, we estimate that 1000 radiation-exposed samples can be analyzed in ~4.11 days with 100 cloned Standard instances. This capacity can be expanded to mitigate backlogs in sample processing. Response time can be further improved by preprocessing samples of known exposures and deriving calibration curves in advance of a radiation emergency.

Security and analytical requirements are fulfilled with independent AWS compute and storage nodes. Data and software are deployed on different platforms that are logically linked to one another during active streaming sessions. Software is maintained in a dormant state until authorized. Images and results are stored in an S3 Bucket accessible only to the user that created them. The AppStream 2.0 environment restricts user uploads to metaphase cell images; outputs and ADCI report files are encrypted. Data are encrypted at rest, in transit between S3 and ADCI Online, and between S3 and the user’s local system.

Users may describe samples that contain personal health information (PHI). PHI is safeguarded in accordance with international data privacy regulations, for example, the Health Insurance Portability and Accountability Act (HIPAA). Compliance is already mandated for sample handling, including interlaboratory transfers^(22)^, management and data processing. Both AppStream 2.0 and S3 storage are HIPAA-eligible services; planned versions of this software are expected to be HIPAA compliant. National or regional data sovereignty requirements are met by co-localizing S3 storage and streaming sessions with user geolocations.

Streaming ADCI Online increases the capacity of locally installed ADCI software to handle radiation incidents of any magnitude. This surge capacity can fulfill burst requirements for analysis of large numbers of samples using parallelized cloud-based instances. Future throughput demands will be met by seamlessly integrating cloud access within the Desktop ADCI software. When configured with an array of GPU-based cloud systems, ADCI Online can estimate radiation doses at population-scale with speeds comparable to either multiple dedicated or supercomputer systems^1^.

## Supporting information

Data Storage and Retrieval web app

## ACKNOWLEDGEMENTS

We are grateful to Drs. Ruth Wilkins and Elizabeth Ainsbury for permitting our reuse of their biodosimetry metaphase image data in this study.

---

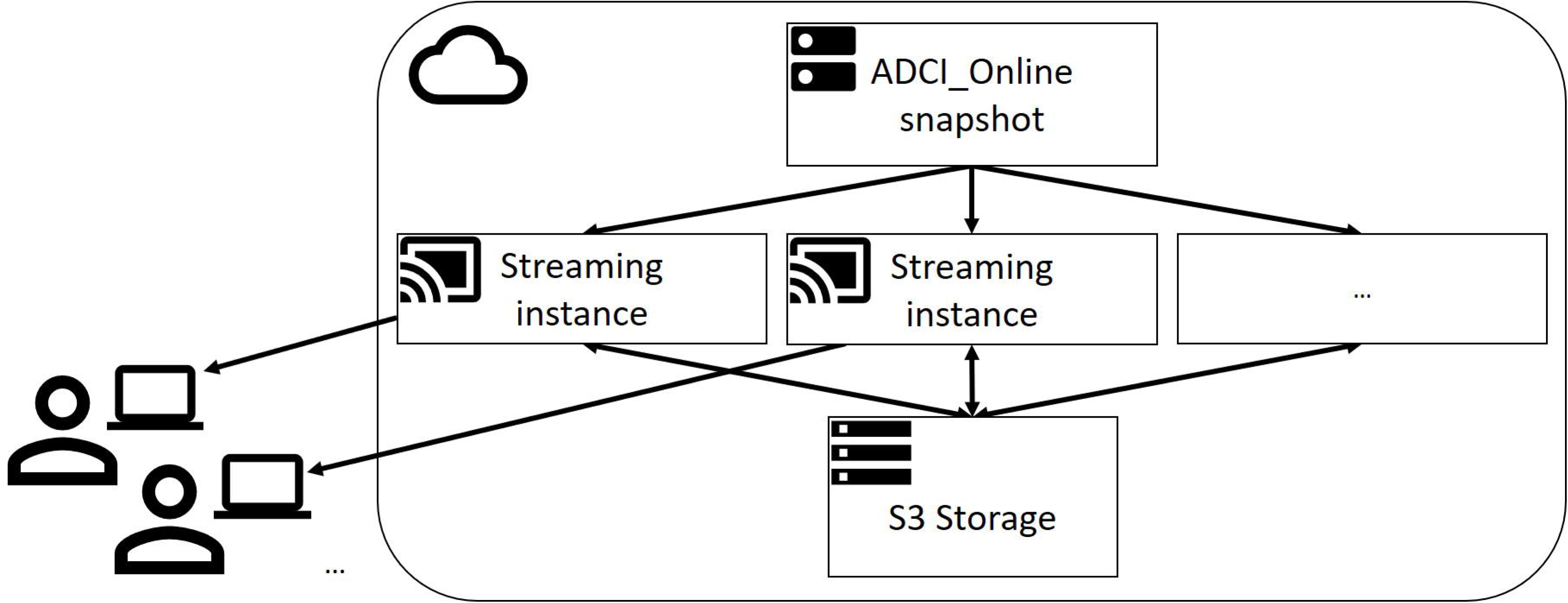

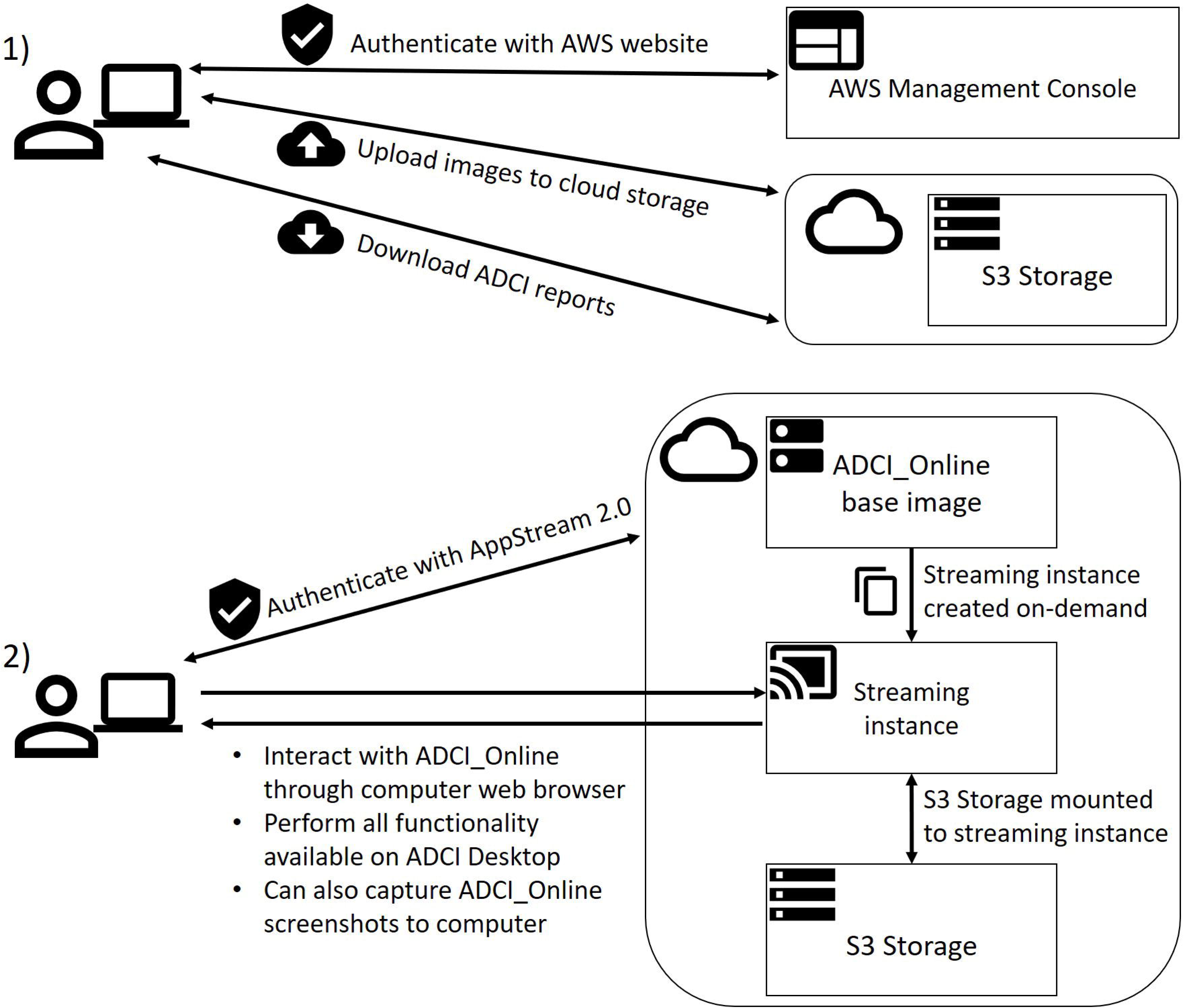

