## Supplementary material for "Radiation Exposure Determination in a Secure, Cloud-Based Online Environment": Data Storage and Retrieval web app: config.js

```
window._config = {
	cognito: {
		USER_POOL_ID: '', //[customize]
		CLIENT_ID: '', //[customize]
		REGION: '', //[customize]
		IDENTITY_POOL_ID: '' //[customize]
	},
	defaults: {
		MAX_FILES_IN_UPLOADED_DIR: 4000,
		MAX_CONCURRENT_UPLOADS: 20,
		SAMPLE_SUBDIR_PREFIX: 'subset',
		BUCKET_NAME: '' //[customize]
	}
}
```
