## Supplementary material for "Radiation Exposure Determination in a Secure, Cloud-Based Online Environment": Data Storage and Retrieval web app: index.php

```
<?php
// Copyright 2021 CytoGnomix Inc. https://www.cytognomix.com
//
// This program is free software: you can redistribute it and/or modify
// it under the terms of the GNU General Public License as published by
// the Free Software Foundation, either version 3 of the License, or
// (at your option) any later version.
//
// This program is distributed in the hope that it will be useful,
// but WITHOUT ANY WARRANTY; without even the implied warranty of
// MERCHANTABILITY or FITNESS FOR A PARTICULAR PURPOSE.  See the
// GNU General Public License for more details.
//
// You should have received a copy of the GNU General Public License
// along with this program.  If not, see <https://www.gnu.org/licenses/>.
?>

<!doctype html>
<html lang="en">
<head>
<meta charset="UTF-8">
<title>ADCI_Online Storage</title>
<link rel="stylesheet" href="https://cdn.jsdelivr.net/npm/bootstrap-icons@1.5.0/font/bootstrap-icons.css">
<link href="https://cdn.jsdelivr.net/npm/bootstrap@5.1.1/dist/css/bootstrap.min.css" rel="stylesheet" integrity="sha384-F3w7mX95PdgyTmZZMECAngseQB83DfGTowi0iMjiWaeVhAn4FJkqJByhZMI3AhiU" crossorigin="anonymous">
<link rel="stylesheet" href="./main.css">
<script src="https://code.jquery.com/jquery-3.6.0.min.js" integrity="sha256-/xUj+3OJU5yExlq6GSYGSHk7tPXikynS7ogEvDej/m4=" crossorigin="anonymous"></script>
<script src="https://cdn.jsdelivr.net/npm/bootstrap@5.1.1/dist/js/bootstrap.bundle.min.js" integrity="sha384-/bQdsTh/da6pkI1MST/rWKFNjaCP5gBSY4sEBT38Q/9RBh9AH40zEOg7Hlq2THRZ" crossorigin="anonymous"></script>
<script src="https://sdk.amazonaws.com/js/aws-sdk-2.977.0.js"></script>

<!-- [customize] Include amazon-cognito-identity, jszip, and FileSaver (not included in Zenodo) -->
<script src="./js/amazon-cognito-identity.js"></script>
<script src="./js/jszip.min.js"></script>
<script src="./js/FileSaver.min.js"></script>
<!-- end [customize] section -->

<script src="./config.js"></script>
<script src="./main.js"></script>
</head>

<body>
	<div class="container">
		<h3 class="mt-3">ADCI_Online Data Storage and Retrieval</h3>
		<p style="margin-bottom:0;font-size:0.9em;">Signing into this resource confirms approval of the terms in this <a target="_blank" href="./documents/ADCI Software_license_updated_10-28-2021_ForWeb.pdf">license</a>.</p><p style="margin-bottom:0;font-size:0.9em;">View <a href="https://adciwiki.cytognomix.com/doku.php?id=adcionline:cloudstorage#signin" target="_blank">sign in documentation</a> on the <a href="https://adciwiki.cytognomix.com/" target="_blank">ADCI Wiki</a>.</p>
		<form>
			<div class="row g-2 mt-0">
				<div class="form-floating col-auto">
					<input type="email" class="form-control" id="inputUserName" name="inputUserName" placeholder="Email" required autofocus>
					<label for="inputUserName">Email address</label>
				</div>
				<div class="form-floating col-auto">
					<input type="password" class="form-control" id="inputPassword" name="inputPassword" placeholder="Password" required>
					<label for="inputPassword">Password</label>
					<p id="forgotPasswordText" style="padding-left:0.5rem;font-size:0.9em;color:#6c757d;text-decoration:underline;cursor:pointer;">Forgot your password?</p>
				</div>
				<div class="col-auto">
					<button class="btn btn-primary" style="height:calc(3.5rem + 2px);" id="signInToCognitoButton" type="button" onclick="signInButton()">Sign in</button>
				</div>			
			</div>	
		</form>


		<div class="card mt-3" id="dirUploadContainer">
			<div class="card-header fw-bold text-white bg-secondary">
				Sample Upload
				<a href="https://adciwiki.cytognomix.com/doku.php?id=adcionline:cloudstorage#sampleupload" target="_blank" class="text-white float-end">View Documentation on the ADCI Wiki</a>
			</div>
			<div class="card-body pb-1 pt-1">
				<p class="section-explanation-text">
					Select a sample (directory) to upload to cloud storage. The selected directory should consist of only image files (tif, tiff, jpg, jpeg, TIF, TIFF, JPG, JPEG) and must not contain any subdirectories. Any files in the sample directory lacking one of these listed file extensions will fail to upload. Samples uploaded to cloud storage are automatically named according to the name of the directory uploaded. For example, if a directory named "Calibration_5Gy" which contains 500 TIFF images is uploaded, the sample will be named "Calibration_5Gy" on cloud storage.
				</p>
				<div class="row mt-3">
					<div class="col-auto">
						<input class="form-control" type="file" id="sample_to_upload" name="sample_to_upload" webkitdirectory mozdirectory />
					</div>
					<div class="col-auto">
						<button class="btn btn-outline-primary" type="button" id="uploadDirToS3">Upload selected sample</button>
					</div>

					<div class="col">
						<div style="font-size:0.9em;" class="row align-items-center float-end">
							<div class="col-auto">
								<label id="batchSizeLabel" style="color:#6c757d;text-decoration:underline;cursor:pointer;">Upload batch size: </label>
							</div>
							<div class="col-auto ps-0">
								<input type="text" style="width:50px;color:#6c757d;cursor:pointer;" id="batchSizeVal" class="form-control-plaintext" readonly value="-">
							</div>			
						</div>
					</div>
				</div>
				
	
				<div class="row mt-3">
					<div class="col">
						<h5>Sample Name</h5>
						<p id="samplename">-</p>
					</div>
					<div class="col">
						<h5>Images in Sample</h5>
						<p id="totalfiles">-</p>
					</div>
					<div class="col">
						<h5>Succeeded</h5>
						<p id="totalsucceeded">-</p>
					</div>
					<div class="col">
						<h5>Failed</h5>
						<p id="totalfailed">-</p>
					</div>
				</div>
				<div class="progress mb-3" style="height: 20px;">
					<div class="progress-bar" role="progressbar" id="diruploadprogressbar" style="width: 0%;" aria-valuenow="0" aria-valuemin="0" aria-valuemax="100">0%</div>
				</div>


				<div class="row mt-3">
					<textarea class="form-control text-dark bg-light" style="font-size:12px;padding-left:calc(var(--bs-gutter-x) * .5);padding-right:calc(var(--bs-gutter-x) * .5);" class="form-control-plaintext" id="consoleOutArea" placeholder="Text output will appear here when uploading a sample" rows="5" readonly></textarea>
				</div>
			</div>
		</div>

		<div class="card mt-3" id="interact_with_uploaded_samples_container">
			<div class="card-header fw-bold text-white bg-secondary">
				Samples on Cloud Storage
				<a href="https://adciwiki.cytognomix.com/doku.php?id=adcionline:cloudstorage#viewsamples" target="_blank" class="text-white float-end">View Documentation on the ADCI Wiki</a>
			</div>
			<div class="card-body pb-1 pt-1">
				<p class="section-explanation-text">
					Samples previously uploaded to cloud storage (using the "Sample Upload" section above) can be viewed here. Click the "Refresh table" button below and the table will be populated with the names of all samples currently stored in the cloud. Click the "Refresh" button under the table heading "Populate/Refresh Image Count" to obtain an image count for the associated sample. This is one method to verify all images in a sample were correctly uploaded. If a sample was uploaded in error or is no longer necessary for your analyses, it can be removed from cloud storage by clicking "Delete" under the table heading "Delete Sample". This table is automatically refreshed when a new sample has been successfully uploaded and when a sample is deleted.
				</p>
				<div class="row mt-3">
					<div>
					<button class="btn btn-outline-primary" type="button" id="listS3BucketButton">Refresh table</button>
					</div>
				</div>
				<div>
					<table class="table table-sm table-striped table-bordered mt-3" id="sample_table">
						<thead>
							<th scope="col">Sample Name</th>
							<th scope="col">Image Count</th>
							<th scope="col">Populate/Refresh Image Count</th>
							<th scope="col">Delete Sample</th>
						</thead>
						<tbody>
							<tr>
								<td>Not yet populated (refresh table)</td>
								<td>-</td>
								<td>-</td>
								<td>-</td>
							</tr>
						</tbody>
					</table>
				</div>
			</div>
		</div>


		<div class="card mt-3" id="download_reports_container">
			<div class="card-header fw-bold text-white bg-secondary">
				Download Reports
				<a href="https://adciwiki.cytognomix.com/doku.php?id=adcionline:cloudstorage#downloadreports" target="_blank" class="text-white float-end">View Documentation on the ADCI Wiki</a>
			</div>
			<div class="card-body pb-1 pt-1">
				<p class="section-explanation-text">
					Reports previously generated by ADCI during an AppStream streaming session can be viewed and downloaded here. Click the "Refresh table" button below and the table will be populated with the names of all ADCI reports currently stored in the cloud. To download a report, click "Download" under the table heading "Download Report". A "Provider report" consists of a single csv file and can be downloaded directly. All other report types consist of an html file and a directory of png image files, thus they are first compressed in zip format before download. The zip process may take several minutes and depends on the number of png files in the report. This table is not automatically refreshed, therefore if you are in an active ADCI_Online streaming session and wish to view recently generated reports, click the "Refresh table" button to obtain a new list of reports.
				</p>
				<div class="row mt-3">
					<div>
						<button class="btn btn-outline-primary" type="button" id="listReportsButton">Refresh table</button>
					</div>
				</div>
				<div>
					<table class="table table-sm table-striped table-bordered mt-3" id="report_table">
						<thead>
							<th scope="col">Report Name</th>
							<th scope="col">Download Report</th>
						</thead>
						<tbody>
							<tr>
								<td>Not yet populated (refresh table)</td>
								<td>-</td>
							</tr>
						</tbody>
					</table>
				</div>
			</div>
		</div>
	</div>

	<div class="text-center p-3 mt-3" style="font-size:0.9em; background-color: rgba(0, 0, 0, 0.05);">
    	Copyright © 2021 <a href="https://cytognomix.com" target="_blank" class="text-reset">CytoGnomix Inc</a>. All rights reserved.
  	</div>

	<iframe id="report_download_iframe" style="display:none;"></iframe>

	<?php
		//modals 
		require_once("./modals/loginrequired.html");
		require_once("./modals/confirmsampledeletion.html");
		require_once("./modals/sampledeletionprogress.html");
		require_once("./modals/newpasswordrequired.html");
		require_once("./modals/forgotpassword.html");
		require_once("./modals/passwordupdated.html");
		require_once("./modals/simplevariablealert.html");
		require_once("./modals/modifybatchsize.html");
	?>

</body>
</html>
```
