## Supplementary material for "Radiation Exposure Determination in a Secure, Cloud-Based Online Environment": Data Storage and Retrieval web app: main.js

```
// Copyright 2021 CytoGnomix Inc. https://www.cytognomix.com
//
// This program is free software: you can redistribute it and/or modify
// it under the terms of the GNU General Public License as published by
// the Free Software Foundation, either version 3 of the License, or
// (at your option) any later version.
//
// This program is distributed in the hope that it will be useful,
// but WITHOUT ANY WARRANTY; without even the implied warranty of
// MERCHANTABILITY or FITNESS FOR A PARTICULAR PURPOSE.  See the
// GNU General Public License for more details.
//
// You should have received a copy of the GNU General Public License
// along with this program.  If not, see <https://www.gnu.org/licenses/>.

// populated upon successfully signing in
var usernameSHA256 = '';
var imagesFolderPrefix = '';
var reportsFolderPrefix = '';

async function signInButton() {
	if(AWS.config.credentials) {
		$('#simpleVariableAlertModal #simpleVariableAlertModal_textContent').html('You are already logged in.');
		$('#simpleVariableAlertModal').modal('show');
		return;
	}	

	var authenticationData = {
		Username : document.getElementById("inputUserName").value,
		Password : document.getElementById("inputPassword").value
	};

	var authenticationDetails = new AmazonCognitoIdentity.AuthenticationDetails(authenticationData);

	var poolData = {
		UserPoolId : _config.cognito.USER_POOL_ID,
		ClientId : _config.cognito.CLIENT_ID
	}
	
	var userPool = new AmazonCognitoIdentity.CognitoUserPool(poolData);

	var userData = {
		Username : document.getElementById("inputUserName").value,
		Pool : userPool
	}

	var cognitoUser = new AmazonCognitoIdentity.CognitoUser(userData);

	var sessionUserAttributes;


	$('#signInToCognitoButton').attr("disabled", true);
	$('#signInToCognitoButton').html('<span class="spinner-grow spinner-grow-sm" role="status" aria-hidden="true"></span> Loading ...');
	$("#signInToCognitoButton").blur();


	cognitoUser.authenticateUser(authenticationDetails, {
		onSuccess: function(result) {
			var emailFromToken = result['idToken']['payload']['email'];
			const sha256Result = digestMessage(emailFromToken);
			sha256Result.then(function(r) {
				usernameSHA256 = JSON.stringify(buf2hex(r)).replaceAll('"', '');
				AWS.config.region = _config.cognito.REGION;
				var cognitoIdpTarget = 'cognito-idp.' + _config.cognito.REGION+'.amazonaws.com/' + _config.cognito.USER_POOL_ID
				AWS.config.credentials = new AWS.CognitoIdentityCredentials({
					IdentityPoolId: _config.cognito.IDENTITY_POOL_ID,
					Logins: {
						[cognitoIdpTarget] : result
							.getIdToken()
							.getJwtToken(),
					},
				});

				AWS.config.credentials.refresh(error => {
					if (error) {
						$('#signInToCognitoButton').attr("disabled", false);
						$('#signInToCognitoButton').html('Sign in');
						AWS.config.credentials = null;
						var loginError = JSON.stringify(error);
						if(error['message'] == "Logins don't match. Please include at least one valid login for this identity or identity pool.") {
							$('#simpleVariableAlertModal #simpleVariableAlertModal_textContent').html('This webpage must be reloaded before you may sign in.');
							$('#simpleVariableAlertModal').modal('show');
						}
						else {
							$('#simpleVariableAlertModal #simpleVariableAlertModal_textContent').html(loginError);
							$('#simpleVariableAlertModal').modal('show');
						}
					} else {
						imagesFolderPrefix = 'user/userpool/' + usernameSHA256 + '/Persistent_ADCI_Data/ADCI_Images/'; //[customize] if path is different
						reportsFolderPrefix = 'user/userpool/' + usernameSHA256 + '/Persistent_ADCI_Data/Reports/'; //[customize] if path is different

						$('#signInToCognitoButton').attr("disabled", true);
						$('#signInToCognitoButton').addClass('btn-outline-success').removeClass('btn-primary');
						$('#signInToCognitoButton').html(' Signed in');	
						$('#signInToCognitoButton').addClass('bi bi-check');
					}
				});

			});


		},
		onFailure: function(err) {
			$('#signInToCognitoButton').attr("disabled", false);
			$('#signInToCognitoButton').html('Sign in');
			
			if (err.code === 'PasswordResetRequiredException') {
				var passwordResetRequiredMsg = 'A password reset is required for this account. Please follow the steps in the &quot;Forgot your password?&quot; link beneath the password input field.';
				$('#simpleVariableAlertModal #simpleVariableAlertModal_textContent').html(passwordResetRequiredMsg);
				$('#simpleVariableAlertModal').modal('show');
			}
			else {
				var loginError = err.message || JSON.stringify(err);
				$('#simpleVariableAlertModal #simpleVariableAlertModal_textContent').html('<p>Error during the login process:</p>' + '<p>' + loginError + '</p>');
				$('#simpleVariableAlertModal').modal('show');
			}


		},
		newPasswordRequired: function(userAttributes, requiredAttributes) {
			// A new password is required if the user was signed up as admin (they always will be in our application)
			$('#signInToCognitoButton').attr("disabled", false);
			$('#signInToCognitoButton').html('Sign in');

			// the api doesn't accept this field back
			delete userAttributes.email_verified;

			// store userAttributes on global variable
			sessionUserAttributes = userAttributes;

			$('#newPasswordRequiredModal #newPasswordRequired_staticEmail').val($('#inputUserName').val());
			$('#newPasswordRequiredModal #newPasswordRequired_updatePasswordButton').attr("disabled", false);
			$('#newPasswordRequiredModal #newPasswordRequired_updatePasswordButton').html('Update password');
			const newPasswordModal = new Promise( function(resolve, reject) {
				$('#newPasswordRequiredModal').modal('show');
				$('#newPasswordRequiredModal #newPasswordRequired_updatePasswordButton').click(function(){
					resolve("Update password");
				});
				$('#newPasswordRequiredModal #newPasswordRequired_cancelButton').click(function(){
					reject("Abandon password update");
				});
			}).then( async function(val) {
				$('#newPasswordRequiredModal #newPasswordRequired_updatePasswordButton').attr("disabled", true);
				$('#newPasswordRequiredModal #newPasswordRequired_updatePasswordButton').html('<span class="spinner-grow spinner-grow-sm" role="status" aria-hidden="true"></span> Loading ...');
				
				var pass1 = $('#newPasswordRequiredModal #newPasswordRequired_password1').val();
				var pass2 = $('#newPasswordRequiredModal #newPasswordRequired_password2').val();
				if(!pass1 || !pass2) {
					$('#newPasswordRequiredModal').modal('hide');
					$('#simpleVariableAlertModal #simpleVariableAlertModal_textContent').html('A new password must be specified in the &quot;New password&quot; field and confirmed in the &quot;Confirm new password&quot; field.');
					$('#simpleVariableAlertModal').modal('show');
					return;
				}

				if(pass1 === pass2) {
					cognitoUser.completeNewPasswordChallenge(pass1, sessionUserAttributes, {
						onSuccess() {
							$('#newPasswordRequiredModal').modal('hide');
							$('#passwordUpdatedModal').modal('show');
							// Clear current credentials in an attempt to avoid requiring a page reload. Users are instructed to log in again after their password has been changed.
							AWS.config.credentials = null;
						},
						onFailure(err) {
							$('#newPasswordRequiredModal').modal('hide');
							var errorMsg = (err.message || JSON.stringify(err));
							$('#simpleVariableAlertModal #simpleVariableAlertModal_textContent').html(errorMsg);
							$('#simpleVariableAlertModal').modal('show');
						},
					}); 
				}
				else {
					$('#newPasswordRequiredModal').modal('hide');
					$('#simpleVariableAlertModal #simpleVariableAlertModal_textContent').html('Passwords do not match.');
					$('#simpleVariableAlertModal').modal('show');
				}
				
				
			}).catch((err) => {
				$('#newPasswordRequiredModal').modal('hide');
				$('#simpleVariableAlertModal #simpleVariableAlertModal_textContent').html(err);
				$('#simpleVariableAlertModal').modal('show');
			});
		}
	});

}

function refreshCredentialsIfNecessary() {
	if (AWS.config.credentials.needsRefresh()) {
		console.log('Credential refresh is necessary');
		AWS.config.credentials.refresh(error => {
			if (error) {
				$('#simpleVariableAlertModal #simpleVariableAlertModal_textContent').html(error);
				$('#simpleVariableAlertModal').modal('show');
			} else {
				console.log('Credentials refreshed');
			}
		});
	}
	else
	{
		console.log('No credential refresh necessary');
	}
}

$(document).on('click', '.report_download_button', async function() {
	var clickedButton = $(this);
	var associatedReportName = $(this).closest('tr').find('.report_name_display').text();
	console.log('Download requested for report: ' + associatedReportName);
	$('.report_download_button, #listReportsButton').attr("disabled", true);
	$(clickedButton).html('<span class="spinner-grow spinner-grow-sm" role="status" aria-hidden="true"></span> Loading ...');
	if(associatedReportName.endsWith('.csv')) {
		await downloadReportSingleCsvFile(associatedReportName);
	}
	else {
		await zipAndDownloadReport(associatedReportName);
	}
	$(clickedButton).html('Download');
	$('.report_download_button, #listReportsButton').attr("disabled", false);
});

$(document).on('click', '.sample_count_button', async function() {
	var clickedButton = $(this);
	var associatedSampleName = $(this).closest('tr').find('.sample_name_display').text();
	var associatedImageCountTd = $(this).closest('tr').find('.sample_image_count_display');
	console.log('Perform file count on sample: ' + associatedSampleName);

	// disable all refresh image count and delete image buttons, set clicked button to display loading message
	$('.sample_count_button, .sample_delete_button, #listS3BucketButton').attr("disabled", true);
	$(clickedButton).html('<span class="spinner-grow spinner-grow-sm" role="status" aria-hidden="true"></span> Loading ...');
	
	const keyCount = await countObjectsInPrefix(associatedSampleName);
	
	// re-enabled all refresh image count and delete image buttons, reset clicked button text to default
	$(clickedButton).html('Refresh');
	$(associatedImageCountTd).html(keyCount);
	$('.sample_count_button, .sample_delete_button, #listS3BucketButton').attr("disabled", false);

});


$(document).ready(function() {
	//apply batch size default value
	$('#batchSizeVal, #modifyBatchSizeModal #modifyBatchSize_currentBatchSizeStatic, #modifyBatchSizeModal #modifyBatchSizeModal_batchSizeSelect').val(_config.defaults.MAX_CONCURRENT_UPLOADS);
	
	$("#listS3BucketButton").click(function(){
		listS3Bucket();
	});
	$("#listReportsButton").click(function(){
		listReportsFolder();
	});
	$("#uploadDirToS3").click(function(){
		uploadDirToS3();
	});
});

$(document).on('click', '#batchSizeLabel, #batchSizeVal', function() {
	$('#modifyBatchSizeModal').modal('show');
});

$(document).on('click', '#modifyBatchSizeModal_updateBatchSizeButton', function() {
	var currentVal = $('#batchSizeVal').val();
	var updatedVal = $('#modifyBatchSizeModal #modifyBatchSizeModal_batchSizeSelect').val();
	if(updatedVal && Number.isInteger(+updatedVal)) {
		if(currentVal === updatedVal) {
			$('#modifyBatchSizeModal #modifyBatchSizeModal_msgArea').html('No change in batch size');
		}
		else
		{
			$('#batchSizeVal, #modifyBatchSizeModal #modifyBatchSize_currentBatchSizeStatic').val(updatedVal);
			$('#modifyBatchSizeModal #modifyBatchSizeModal_msgArea').html('Batch size successfully updated');
		}		
	}
	else {
		$('#modifyBatchSizeModal #modifyBatchSizeModal_msgArea').html('Specified batch size is not a number. Batch size not updated.');
	}
});


$(document).on('click', '.sample_delete_button', function() {
	if(AWS.config.credentials == null) {
		$('#loginRequiredModal').modal('show');
		return;
	}

	var associatedSampleName = $(this).closest('tr').find('.sample_name_display').text();
	console.log('Delete the sample: ' + associatedSampleName);

	$('#confirmSampleDeletionModal .confirmSampleDeleteModal-SampleName').html(associatedSampleName);
	$('#confirmSampleDeletionModal').modal('show');
});

$(document).on('click', '#sampleDeletionConfirmedButton', async function() {
	var sampleToBeDeleted = $('#confirmSampleDeletionModal .confirmSampleDeleteModal-SampleName').first().html();
	$('#confirmSampleDeletionModal').modal('hide');
	$('#confirmSampleDeletionModal .confirmSampleDeleteModal-SampleName').html('UNDEFINED');
	
	await deleteSample(sampleToBeDeleted);	
});


$(document).on('click', '#forgotPasswordText', function() {
	$('#forgotPasswordModal').modal('show');
});

$(document).on('click', '#forgotPassword_sendCodeButton', function() {
	var forgotPasswordPoolData = {
		UserPoolId : _config.cognito.USER_POOL_ID,
		ClientId : _config.cognito.CLIENT_ID
	}
	
	var forgotPasswordUserPool = new AmazonCognitoIdentity.CognitoUserPool(forgotPasswordPoolData);
	var forgotPasswordUserData = {
		Username : document.getElementById("forgotPassword_userName").value,
		Pool : forgotPasswordUserPool
	}

	var forgotPasswordUser = new AmazonCognitoIdentity.CognitoUser(forgotPasswordUserData);
	$('#forgotPassword_sendCodeResponseArea').html('<div class="text-center"><div class="spinner-grow text-secondary" role="status"><span class="visually-hidden"></span></div></div>');

	forgotPasswordUser.forgotPassword({
		onSuccess: function(data) {
			$('#forgotPassword_sendCodeResponseArea').html('Code sent successfully');
		},
		onFailure: function(err) {
			var sendCodeResponse = err.message || JSON.stringify(err);
			$('#forgotPassword_sendCodeResponseArea').html(sendCodeResponse); 
			return;
		}
	});
});

$(document).on('click', '#forgotPassword_updatePasswordButton', function() {

	var forgotPasswordPoolData = {
		UserPoolId : _config.cognito.USER_POOL_ID,
		ClientId : _config.cognito.CLIENT_ID
	}
	
	var forgotPasswordUserPool = new AmazonCognitoIdentity.CognitoUserPool(forgotPasswordPoolData);
	var forgotPasswordUserData = {
		Username : document.getElementById("forgotPassword_userName").value,
		Pool : forgotPasswordUserPool
	}
	
	var forgotPasswordUser = new AmazonCognitoIdentity.CognitoUser(forgotPasswordUserData);

	var verificationCode = $('#forgotPasswordModal #forgotPassword_code').val();
	var pass1 = $('#forgotPasswordModal #forgotPassword_password1').val();
	var pass2 = $('#forgotPasswordModal #forgotPassword_password2').val();
	
	if(!forgotPasswordUserData.Username) {
		$('#forgotPassword_errorArea').html('An email address must be specified.');
		return;
	}
	if(!verificationCode) {
		$('#forgotPassword_errorArea').html('A verification code must be specified.');
		return;
	}
	if(!pass1 || !pass2) {
		$('#forgotPassword_errorArea').html('A new password must be specified in the &quot;New password&quot; field and confirmed in the &quot;Confirm new password&quot; field.');
		return;
	}


	if(pass1 !== pass2) {
		$('#forgotPassword_errorArea').html('The input passwords do not match.');
		return;
	}
	else {
		$('#forgotPassword_errorArea').html('<div class="text-center"><div class="spinner-grow text-secondary" role="status"><span class="visually-hidden"></span></div></div>');
		forgotPasswordUser.confirmPassword(verificationCode, pass1, {
			onSuccess() {
				$('#forgotPassword_errorArea').html('New password successfully applied. Please log in again.');
				$('#forgotPasswordModal').modal('hide');
				$('#passwordUpdatedModal').modal('show');
			},
			onFailure(err) {
				var updatePasswordResponse = err.message || JSON.stringify(err);
				$('#forgotPassword_errorArea').html(updatePasswordResponse);
			},
		});
	}
});


async function zipAndDownloadReport(reportName) {
	reportsFolder = 'user/userpool/' + usernameSHA256 + '/Persistent_ADCI_Data/Reports/';
	targetReportKey = reportsFolder + reportName + '/';

	var s3 = new AWS.S3({apiVersion: "2006-03-01", params:{Bucket: _config.defaults.BUCKET_NAME}});
	const listObjectsResults = await getAllKeysAtPrefix(s3, targetReportKey);

	var zipper = new JSZip();
	var errorCaught = false;
	for(var i = 0; i < listObjectsResults.keys.length; ++i) {
		//skip 'folders' in S3
		if(listObjectsResults.keys[i].endsWith('/')) {
			continue;
		}
		paramsForGetObject = {
			Bucket: _config.defaults.BUCKET_NAME,
			Key: listObjectsResults.keys[i]
		}
		try {
			const getObjResponse = await s3.getObject(paramsForGetObject).promise();
			var blob = new Blob([getObjResponse.Body], {type: getObjResponse.ContentType});
			var trimmedKey = listObjectsResults.keys[i].replace(targetReportKey, '');
			zipper.file(trimmedKey, blob);
		}
		catch (ex) {
			console.log('Caught error while performing getObject:');
			console.log(ex);
			$('#simpleVariableAlertModal #simpleVariableAlertModal_textContent').html('<p>Error while attempting to download report:</p>' + '<p>' + ex.message + '</p>');
			$('#simpleVariableAlertModal').modal('show');
			errorCaught = true;
			break;
		}
	}

	if(!errorCaught) {
		await zipper.generateAsync({type : "blob"}).then(function(blob) {
			var zipFileName = reportName + '.zip';
			window.saveAs(blob, zipFileName);
		}, function(err) {
			$('#simpleVariableAlertModal #simpleVariableAlertModal_textContent').html(err);
			$('#simpleVariableAlertModal').modal('show');
		});
	}
}


async function downloadReportSingleCsvFile(reportName) {
	reportsFolder = 'user/userpool/' + usernameSHA256 + '/Persistent_ADCI_Data/Reports/';
	targetReportKey = reportsFolder + reportName;

	var s3 = new AWS.S3({apiVersion: "2006-03-01", params:{Bucket: _config.defaults.BUCKET_NAME}});
	paramsForGetObject = {
		Bucket: _config.defaults.BUCKET_NAME,
		Key: targetReportKey
	}
	try {
		const getObjResponse = await s3.getObject(paramsForGetObject).promise();
		var blob = new Blob([getObjResponse.Body], {type: "text/csv;charset=utf-8"});
		var trimmedKey = targetReportKey.replace(reportsFolder, '');
		window.saveAs(blob, trimmedKey);
	}
	catch (ex) {
		console.log('Caught error while performing getObject:');
		console.log(ex);
		$('#simpleVariableAlertModal #simpleVariableAlertModal_textContent').html('<p>Error while attempting to download report:</p>' + '<p>' + ex.message + '</p>');
		$('#simpleVariableAlertModal').modal('show');
	}
}


async function deleteSample(sampleToBeDeleted) {
	// clear and configure sample deletion modal
	updateSampleDeletionProgressBar(0, 1);
	$('#sampledeletion_samplename').html(sampleToBeDeleted);
	$('#sampledeletion_totalfiles').html('<div class="spinner-border text-secondary" role="status"><span class="visually-hidden">Loading...</span></div>');
	$('#sampledeletion_totaldeleted').html('-');
	$('#closeSampleDeletionModalButton').attr("disabled", true);

	// show sample deletion modal, prevent modal from closing when clicking outside modal area or pressing escape
	$('#sampleDeletionProgressModal').modal('show');

	// get list of keys in sample
	var sampleFolder = imagesFolderPrefix + sampleToBeDeleted + '/';
	var s3 = new AWS.S3({apiVersion: "2006-03-01", params:{Bucket: _config.defaults.BUCKET_NAME}});
	console.log('Sample key: ' + sampleFolder)
	try {
		const listObjectsResults = await getAllKeysAtPrefix(s3, sampleFolder);

		//stop spinner on image count area and display obtained image count
		var totalFilesInSample = listObjectsResults.keys.length;
		var totalSuccessfullyDeleted = 0;
		$('#sampledeletion_totalfiles').html(totalFilesInSample);

		// delete sample in chunks of 1000 images (max allowed)
		var chunkSize = 1000;
		for (var i = 0; i < totalFilesInSample; i += chunkSize) {
			var arrChunk = listObjectsResults.keys.slice(i, i + chunkSize);
			var objectsToDelete = {Objects: []};
			for(var j = 0; j < arrChunk.length; j++) {
				objectsToDelete.Objects.push({Key: arrChunk[j]});
			}
			var params = {Bucket: _config.defaults.BUCKET_NAME, Delete: objectsToDelete};
			
			const deleteResponse = await s3.deleteObjects(params).promise();
			if(deleteResponse.Deleted) {
				var successfullyDeleted = deleteResponse.Deleted.length;
				totalSuccessfullyDeleted += successfullyDeleted;
				updateSampleDeletionProgressBar(totalSuccessfullyDeleted, totalFilesInSample);
				$('#sampledeletion_totaldeleted').html(totalSuccessfullyDeleted);
			}
		}
	}
	catch(e) {
		$('#sampledeletion_errorarea').html('An error occurred: ' + e.message);
		// if spinner is still present, clear it
		if($('#sampledeletion_totalfiles > div').hasClass('spinner-border')) {
			$('#sampledeletion_totalfiles').html('Error');
		}
	}
	finally {
		$('#closeSampleDeletionModalButton').attr("disabled", false);
		//refresh table after sample deletion (or attempted deletion)
		listS3Bucket();
	}	
}

async function getAllKeysAtPrefix(s3, targetPrefix) {
	var listObjectsResults = {keys: [], folders: []};

	var loopAgain = false;
	var nextContinuationToken = '';
	do {
		var listParams = {Prefix: targetPrefix, Delimiter: '/', MaxKeys: 1000};
		if(nextContinuationToken) {
			Object.assign(listParams, {ContinuationToken: nextContinuationToken});
		}
		loopAgain = false;
		nextContinuationToken = '';

		const listResponse = await s3.listObjectsV2(listParams).promise();
		listResponse.Contents.forEach((element, index) => {
			listObjectsResults.keys.push(element.Key);
		});
		listResponse.CommonPrefixes.forEach((element, index) => {
			listObjectsResults.folders.push(element.Prefix);	
		});

		if(listResponse.IsTruncated === true && listResponse.NextContinuationToken) {
			nextContinuationToken = listResponse.NextContinuationToken;
			loopAgain = true;
		}
	}
	while(loopAgain === true);


	//if there were any subfolders
	for (var i = 0; i < listObjectsResults.folders.length; i++) {
		var loopSubfolderAgain = false;
		var nextContinuationTokenForSubfolder = '';
		do {
			var listParams = {Prefix: listObjectsResults.folders[i], Delimiter: '/', MaxKeys: 1000};
			if(nextContinuationTokenForSubfolder) {
				Object.assign(listParams, {ContinuationToken: nextContinuationTokenForSubfolder});
			}
			loopSubfolderAgain = false;
			nextContinuationTokenForSubfolder = '';

			const listResponse = await s3.listObjectsV2(listParams).promise();
			listResponse.Contents.forEach((element) => {
				listObjectsResults.keys.push(element.Key);
			});
			listResponse.CommonPrefixes.forEach((element) => {
				listObjectsResults.folders.push(element.Prefix);	
			});

			if(listResponse.IsTruncated === true && listResponse.NextContinuationToken) {
				nextContinuationTokenForSubfolder = listResponse.NextContinuationToken;
				loopSubfolderAgain = true;
			}
		}
		while(loopSubfolderAgain === true);
	}

	return listObjectsResults;
}


async function countObjectsInPrefix(targetSampleName) {
	if(AWS.config.credentials == null) {
		$('#loginRequiredModal').modal('show');
		return;
	}

	var sampleFolder = imagesFolderPrefix + targetSampleName + '/';
	var s3 = new AWS.S3({apiVersion: "2006-03-01", params:{Bucket: _config.defaults.BUCKET_NAME}});

	const listObjectsResults = await getAllKeysAtPrefix(s3, sampleFolder);
	
	return listObjectsResults.keys.length;
}

function listS3Bucket() {	
	if(AWS.config.credentials == null) {
		$('#loginRequiredModal').modal('show');
		return;
	}


	refreshCredentialsIfNecessary();
	if (! AWS.config.credentials.needsRefresh()) {
		var s3 = new AWS.S3({apiVersion: "2006-03-01", params:{Bucket: _config.defaults.BUCKET_NAME}});

		
		$('#listS3BucketButton, .sample_count_button, .sample_delete_button').attr("disabled", true);
		$('#listS3BucketButton').html('<span class="spinner-grow spinner-grow-sm" role="status" aria-hidden="true"></span> Loading ...');
		

		s3.listObjects({ Prefix: imagesFolderPrefix, Delimiter: "/" }, function(err, data) {
			$('#listS3BucketButton, .sample_count_button, .sample_delete_button').attr("disabled", false);
			$('#listS3BucketButton').html('Refresh table');
			if (err) {
				$('#simpleVariableAlertModal #simpleVariableAlertModal_textContent').html(err);
				$('#simpleVariableAlertModal').modal('show');
			}
			else {
				$('#sample_table > tbody').html('');

				if(data.CommonPrefixes.length === 0) {
					var rowToAppend = '<tr><td>No content</td><td>-</td><td>-</td><td>-</td></tr>';
					$('#sample_table tbody').append(rowToAppend);
					return;
				}

				data.CommonPrefixes.forEach((element, index) => {
					var fullKeyForCommonPrefix = element.Prefix;
					var sampleNameOnly = fullKeyForCommonPrefix.replace(imagesFolderPrefix, '');
					//strip trailing slash
					sampleNameOnly = sampleNameOnly.replace(/\/$/, '');
					var rowToAppend = 	'<tr>\
											<td class="sample_name_display">'+sampleNameOnly+'</td>\
											<td class="sample_image_count_display">-</td>\
											<td>\
												<button class="btn btn-outline-primary sample_count_button" type="button">Refresh</button>\
											</td>\
											<td>\
												<button class="btn btn-outline-danger sample_delete_button" type="button">Delete</button>\
											</td>\
										</tr>';
					$('#sample_table tbody').append(rowToAppend);
				});

			}
		});
	}
}

async function listReportsFolder() {	
	if(AWS.config.credentials == null) {
		$('#loginRequiredModal').modal('show');
		return;
	}

	refreshCredentialsIfNecessary();
	if (! AWS.config.credentials.needsRefresh()) {
		var s3 = new AWS.S3({apiVersion: "2006-03-01", params:{Bucket: _config.defaults.BUCKET_NAME}});

		$('.report_download_button, #listReportsButton').attr("disabled", true);
		$('#listReportsButton').html('<span class="spinner-grow spinner-grow-sm" role="status" aria-hidden="true"></span> Loading ...');		
		s3.listObjectsV2({ Prefix: reportsFolderPrefix, Delimiter: "/" }, function(err, data) {
			$('.report_download_button, #listReportsButton').attr("disabled", false);
			$('#listReportsButton').html('Refresh table');
			if (err) {
				$('#simpleVariableAlertModal #simpleVariableAlertModal_textContent').html(err);
				$('#simpleVariableAlertModal').modal('show');
			}
			else {
				$('#report_table > tbody').html('');
				
				//since the downloads folder itself is returned, reports are present if there are 2 or more keys
				if(data.CommonPrefixes.length === 0) {
					var rowToAppend = '<tr><td>No content</td><td>-</td></tr>';
					$('#report_table tbody').append(rowToAppend);
					return;
				}
				
				var reportNames = [];
				data.CommonPrefixes.forEach((element) => {
					var fullKeyForCommonPrefix = element.Prefix;
					var reportNameOnly = fullKeyForCommonPrefix.replace(reportsFolderPrefix, '');
					//strip trailing slash
					reportNameOnly = reportNameOnly.replace(/\/$/, '');
					reportNames.push(reportNameOnly);
				});

				data.Contents.forEach((element) => {
					if(element.Size > 0) {
						var fullKeyForCommonPrefix = element.Key;
						var reportNameOnly = fullKeyForCommonPrefix.replace(reportsFolderPrefix, '');
						reportNames.push(reportNameOnly);
					}
				});

				reportNames.forEach((element) => {
					var rowToAppend = 	'<tr>\
											<td class="report_name_display">'+element+'</td>\
											<td>\
												<button class="btn btn-outline-primary report_download_button" type="button">Download</button>\
											</td>\
										</tr>';
					$('#report_table tbody').append(rowToAppend);
				});

			}
		});
	}
}

async function digestMessage(message) {
	const encoder = new TextEncoder();
	const data = encoder.encode(message);
	const hash = await crypto.subtle.digest('SHA-256', data);
	return hash;
  }

function buf2hex(buffer) {
    return Array.prototype.map.call(new Uint8Array(buffer), x => ('00' + x.toString(16)).slice(-2)).join('');
}


function sleep(ms) {
	return new Promise(resolve => setTimeout(resolve, ms));
  }

function writeToConsole(text, addNewline) {
	if(addNewline) {
		text += '\n';
	}
	var targetTextArea = document.getElementById('consoleOutArea');
	targetTextArea.value += text;
	targetTextArea.scrollTop = targetTextArea.scrollHeight;
}

function updateSampleDeletionProgressBar(complete, total) {
	var percentComplete = (complete / total) * 100;	
	percentComplete = Math.round(percentComplete * 100) / 100
	$('#sampledeletion_progressbar').css('width', percentComplete+'%').attr('aria-valuenow', percentComplete);
	$('#sampledeletion_progressbar').text(percentComplete + '%');
}

function updateProgressBar(complete, total) {
	$('#diruploadprogressbar').removeClass('bg-danger');
	var percentComplete = (complete / total) * 100;	
	percentComplete = Math.round(percentComplete * 100) / 100
	$('#diruploadprogressbar').css('width', percentComplete+'%').attr('aria-valuenow', percentComplete);
	$('#diruploadprogressbar').text(percentComplete + '%');
}

function setProgressBarToCancelledState() {
	$('#diruploadprogressbar').css('width', '100%').attr('aria-valuenow', '100');
	$('#diruploadprogressbar').addClass('bg-danger');
	$('#diruploadprogressbar').text('Upload Cancelled');
}

async function uploadDirToS3() {
	if(AWS.config.credentials == null) {
		$('#loginRequiredModal').modal('show');
		return;
	}

	var beginTime = performance.now();
	var batchSize = $('#batchSizeVal').val();
	var filesInDir = document.getElementById("sample_to_upload").files;
	updateProgressBar(0, filesInDir.length);

	if(filesInDir.length == 0) {
		writeToConsole('No sample directory specified or no files are present in specified directory. Cancelling upload ...', true);
		return;
	}

	$('#uploadDirToS3, #sample_to_upload').attr("disabled", true);
	$('#uploadDirToS3').html('<span class="spinner-grow spinner-grow-sm" role="status" aria-hidden="true"></span> Uploading ...');

	var adciImagesPrefix = imagesFolderPrefix;

	var sampleMustBeSplitIntoSubdirectories = false;
	if(filesInDir.length > _config.defaults.MAX_FILES_IN_UPLOADED_DIR) {
		sampleMustBeSplitIntoSubdirectories = true;
	}

	var totalAcceptableFilesInDir = 0;
	var sampleNameToUpload = '';
	for(i = 0; i < filesInDir.length; i++) {
		if (filesInDir[i].webkitRelativePath.match(/\//g).length == 1) {
			totalAcceptableFilesInDir++;
			if(sampleNameToUpload == '') {
				var splitRelativePath = filesInDir[i].webkitRelativePath.toString().split('/');
				sampleNameToUpload = splitRelativePath[0]
				$('#samplename').text(sampleNameToUpload);
				sampleNameDisplaySet = true;
			}
			
		}
	}
	
	$('#totalfiles').text(totalAcceptableFilesInDir);

	writeToConsole('=== Sample Name: ' + sampleNameToUpload + ' ===', true);
	writeToConsole('Base image directory on S3: ' + adciImagesPrefix, true);
	writeToConsole('Base upload directory for this sample: ' + adciImagesPrefix + sampleNameToUpload, true);

	var totalSucceeded = 0;
	var totalFailed = 0;
	var uploadParams = [];
	var counterOfFilesInSpecifiedDirectory = 0;
	var i = 0;

	var uploadCancelled = false;
	while (i < filesInDir.length) {
		if (filesInDir[i].webkitRelativePath.match(/\//g).length == 1) {
			
			targetKey = '';
			var splitRelativePath = filesInDir[i].webkitRelativePath.toString().split('/');
			if(sampleMustBeSplitIntoSubdirectories) {
				var targetSubdirNumber = Math.floor( counterOfFilesInSpecifiedDirectory / _config.defaults.MAX_FILES_IN_UPLOADED_DIR ) + 1;
				var targetSubdir = _config.defaults.SAMPLE_SUBDIR_PREFIX + targetSubdirNumber.toString();
				targetKey = adciImagesPrefix + splitRelativePath[0] + '/' + targetSubdir + '/' + splitRelativePath[1];
			}	
			else {
				targetKey = adciImagesPrefix + filesInDir[i].webkitRelativePath;
			}
			
			uploadParams.push( {Bucket: _config.defaults.BUCKET_NAME, Key: targetKey, Body: filesInDir[i]} );

			if(uploadParams.length > 0) {
				if(uploadParams.length >= batchSize || i + 1 == filesInDir.length) {
					var promises = [];
					uploadParams.forEach((element, index) => {
						var upload = new AWS.S3.ManagedUpload({params: uploadParams[index]});
						promises.push( upload.promise() );
					})
					var currentBatchUploadsSucceeded = 0;
					var currentBatchUploadsFailed = 0;
					await Promise.allSettled(promises).then((respArr) => {
						respArr.forEach((element, index) => {
							if(element.status == 'fulfilled') {
								currentBatchUploadsSucceeded++;
							}
							else {
								var imageNameOnly = uploadParams[index].Key.replace(imagesFolderPrefix + sampleNameToUpload + '/', '');
								writeToConsole(imageNameOnly + ' - Upload failed. Reason: ' + element.reason, true);
								currentBatchUploadsFailed++;
							}
						})
					});
					if(currentBatchUploadsFailed > 0) {
						writeToConsole(currentBatchUploadsFailed + '/' + uploadParams.length + ' files in upload batch failed to upload.', true);
					}
					
					totalSucceeded += currentBatchUploadsSucceeded;
					totalFailed += currentBatchUploadsFailed;
					$('#totalsucceeded').text(totalSucceeded);
					$('#totalfailed').text(totalFailed);
					var totalProcessed = totalSucceeded + totalFailed;
					updateProgressBar(totalProcessed, totalAcceptableFilesInDir);
					uploadParams = [];
					if(totalFailed > totalSucceeded) {
						uploadCancelled = true;
						var uploadCancelMsg = 'Upload failures are too frequent. Cancelling upload.'
						writeToConsole(uploadCancelMsg, true);
						break;
					}
				}
			}
			counterOfFilesInSpecifiedDirectory++;
		}
		i++;
	}
	var endTime = performance.now();
	var elapsedTimeInMinutes = (endTime - beginTime) / 60000;
	elapsedTimeInMinutes = Math.round(elapsedTimeInMinutes * 100) / 100
	if(uploadCancelled) {
		writeToConsole('Sample upload cancelled after: ' + elapsedTimeInMinutes + ' minutes.', true);
		setProgressBarToCancelledState();
	}
	else {
		writeToConsole('Sample uploaded in: ' + elapsedTimeInMinutes + ' minutes.', true);
	}
	$('#uploadDirToS3, #sample_to_upload').attr("disabled", false);
	$('#uploadDirToS3').html('Upload selected sample');
}
```
