## Supplementary material for "Radiation Exposure Determination in a Secure, Cloud-Based Online Environment": Data Storage and Retrieval web app: forgotpassword.html

##### Reset Password

First, request that a verification code be sent to the e-mail address you use to login to ADCI\_Online. After you have received the code, enter it along with a new password of your choice and click "Update password". Verification codes are valid for 24 hours.

Please note: Passwords must be at least 8 characters long and require at least one number, special character, uppercase letter, and lowercase letter.

Email:

Send code

Code:

New password:

Confirm new password:

Cancel
Update password
