## Supplementary material for "Radiation Exposure Determination in a Secure, Cloud-Based Online Environment": Data Storage and Retrieval web app: modifybatchsize.html

##### Modify Batch Size

To improve upload speeds, batches of metaphase images are uploaded to cloud storage concurrently. All images in a batch are uploaded simultaneously, then another batch is created and uploaded. This process repeats until all metaphase images in a sample have been uploaded.

Selection of an appropriate batch size is a balance between overall upload speed and the load higher batch sizes place on the system performing the upload. Factors to consider when selecting a batch size include 1) CPU performance, 2) disk read speed, and 3) internet bandwidth. When attempting to maximize upload speed, the goal is to increase batch size to utilize all available internet bandwidth. However, high batch sizes may slow a computing system and cause it to become less responsive. If the system cannot keep up with the demand, increased batch size will not offer any improvement. The dropdown box below contains recommended batch sizes for a variety of systems.

Current batch size:

Select batch size:

2 - debug
10 - Laptop using a wireless internet connection
20 - Standard workstation
30 - High performance desktop system, wired internet access
50 - Superior performance desktop system paired with high throughput internet access

Close
Update batch size
