## Supplementary material for "Radiation Exposure Determination in a Secure, Cloud-Based Online Environment": Data Storage and Retrieval web app: newpasswordrequired.html

##### New Password Required

The password you entered is a temporary password created automatically and must be updated. Please enter a new password below.

Please note: Passwords must be at least 8 characters long and require at least one number, special character, uppercase letter, and lowercase letter.

Email:

New
password:

Confirm new
password:

Cancel
Update password
