## Supplementary Document for "Radiation Exposure Determination in a Secure, Cloud-Based Online Environment"

### Companion Screenshots for: Radiation Exposure Determination in a Secure, Cloud-based Online Environment

### Upload Samples to Encrypted Cloud Storage

<u>Time Required:</u>
~1 min per 1000 metaphase images*

54,595 metaphase images (41.7GB on disk) uploaded in 53.64 min.

- Users sign in to our Javascript web app that provides an interface to cloud storage.
- Sign in credentials are provided in an e-mail sent to the user by system administrators.

- Users browse their local system for a directory containing a sample to upload and select "Upload Selected Sample".
- All metaphase images associated with a sample must be present in a single directory.
- The percentage of metaphase images uploaded is displayed using a progress bar, and the number of metaphase images successfully uploaded is displayed.
- Upon completion of the upload, the time required to perform it is displayed in the output console.
- This process is repeated for each sample to be uploaded.

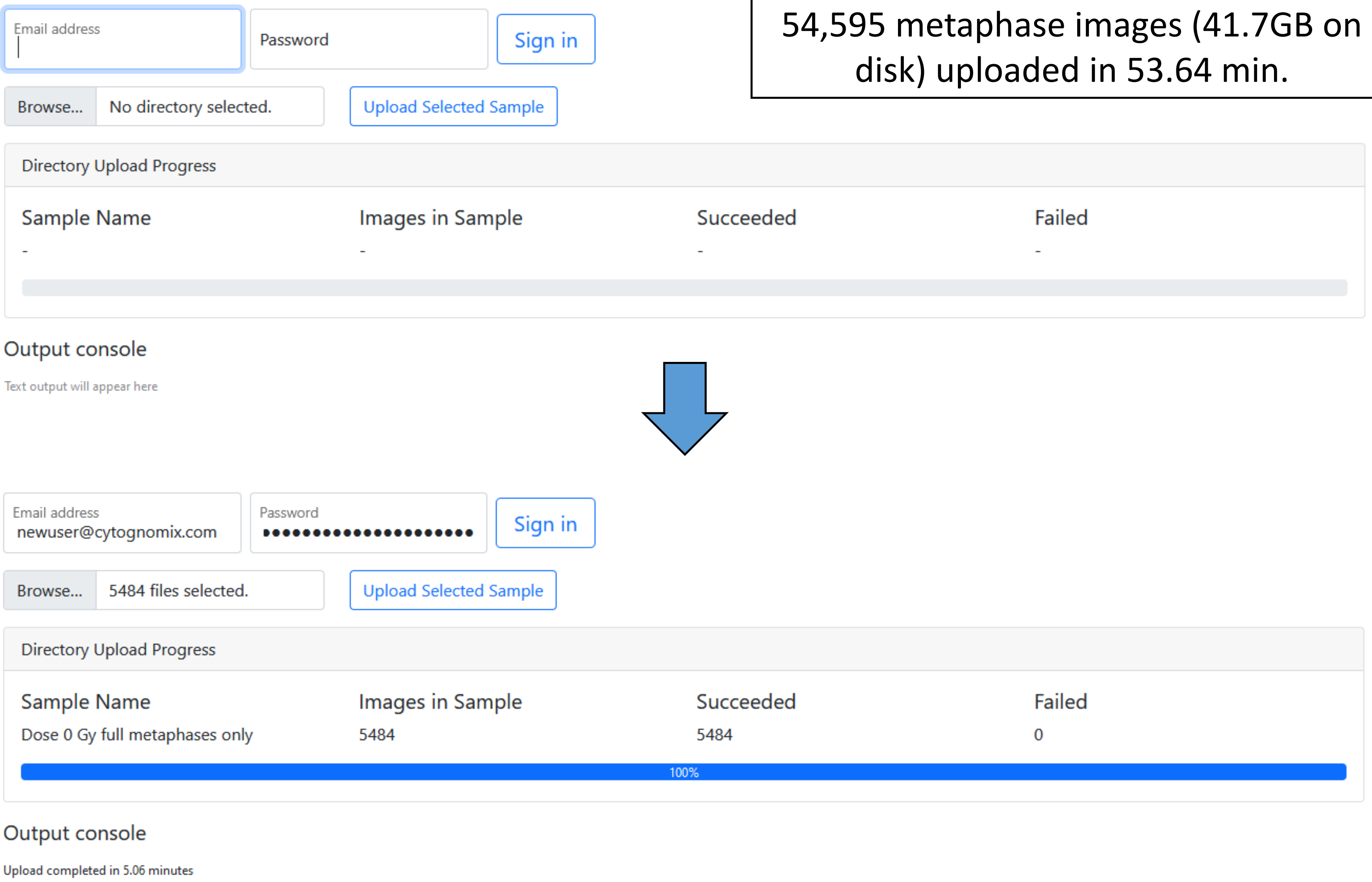


* Dependent on internet bandwidth and number of parallel uploads (also dependent on CPU and disk read speed).

### Access ADCI_Online

Time Required:
< 5 minutes

- Users access ADCI_Online through AWS AppStream 2.0.
- Sign in credentials are provided in an e-mail sent by AWS.
- AppStream is accessible through a web browser (recommended) or a Windows client installed on the local system.

- “Desktop” or “ADCI” stream views are available. “Desktop” presents a Windows desktop from which ADCI can be executed, the “ADCI” view consists only of the ADCI software.

- A new streaming session is created in approximately two minutes.

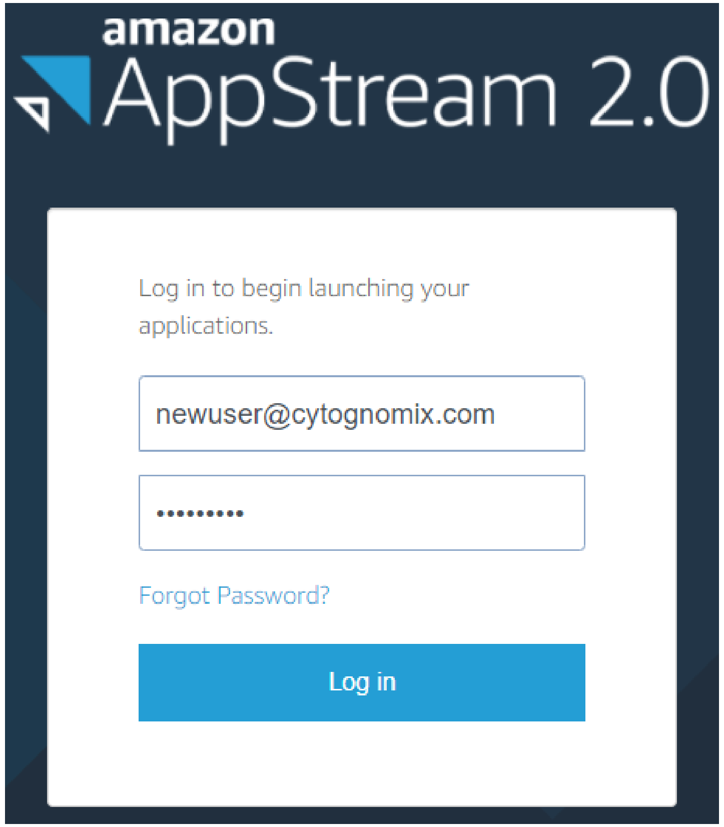


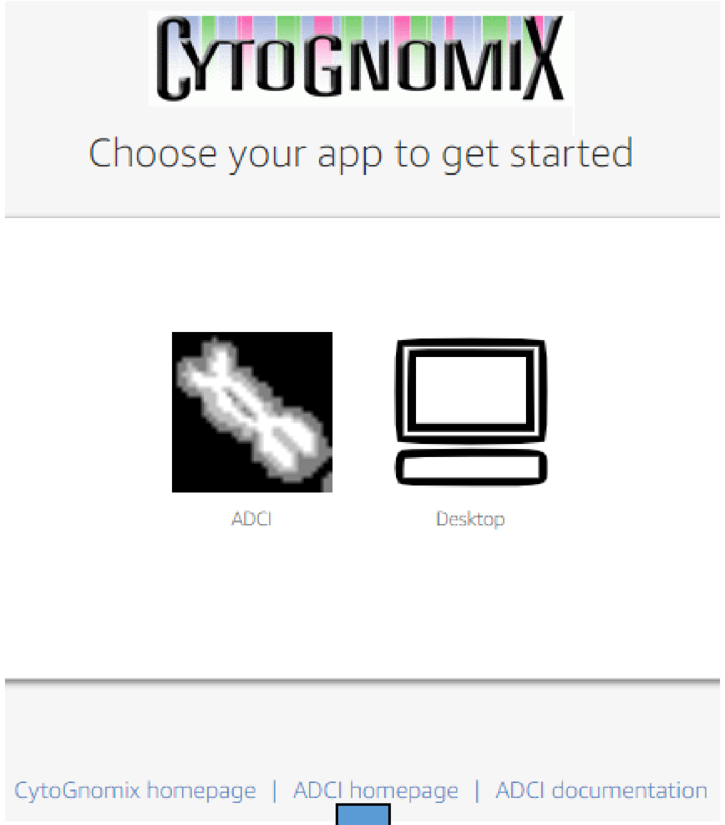


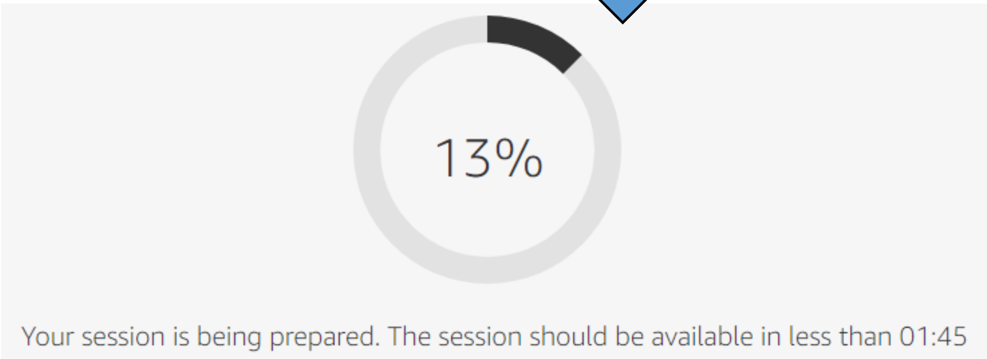

### Access ADCI_Online (continued)

Time Required:
< 10 seconds

- ADCI is now executing on the cloud.
- Cloud storage has been automatically mounted to the streaming instance, granting access to the previously uploaded samples.

- Subsequent steps are similar to those performed on a standard desktop implementation of ADCI installed locally, thus the ADCI manual (https://adciwiki.cytognomix.com) can be consulted for additional details.

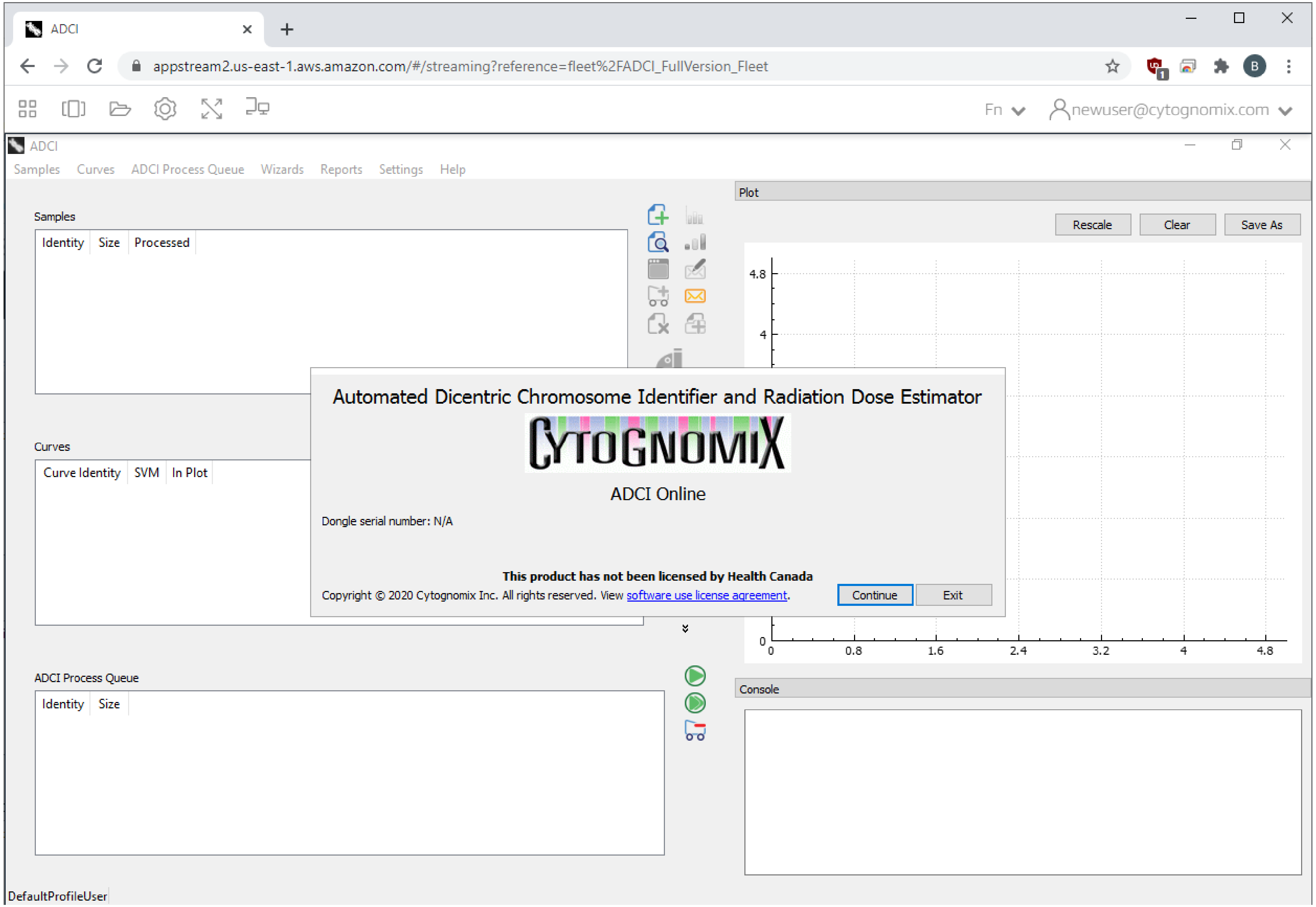

### Create New Samples

Time Required:
< 10 minutes

- To add a sample to ADCI, a directory containing metaphase images must be selected.
- When browsing the streaming instance for a directory, the "PERSISTENT_ADCI_DATA" directory will be displayed by default. The subdirectory "ADCI_Images" contains samples uploaded using the Javascript web app.
- A unique ID for the sample must be specified. The name of the sample directory can be used if desired.
- This process is repeated for all samples to be processed by ADCI.

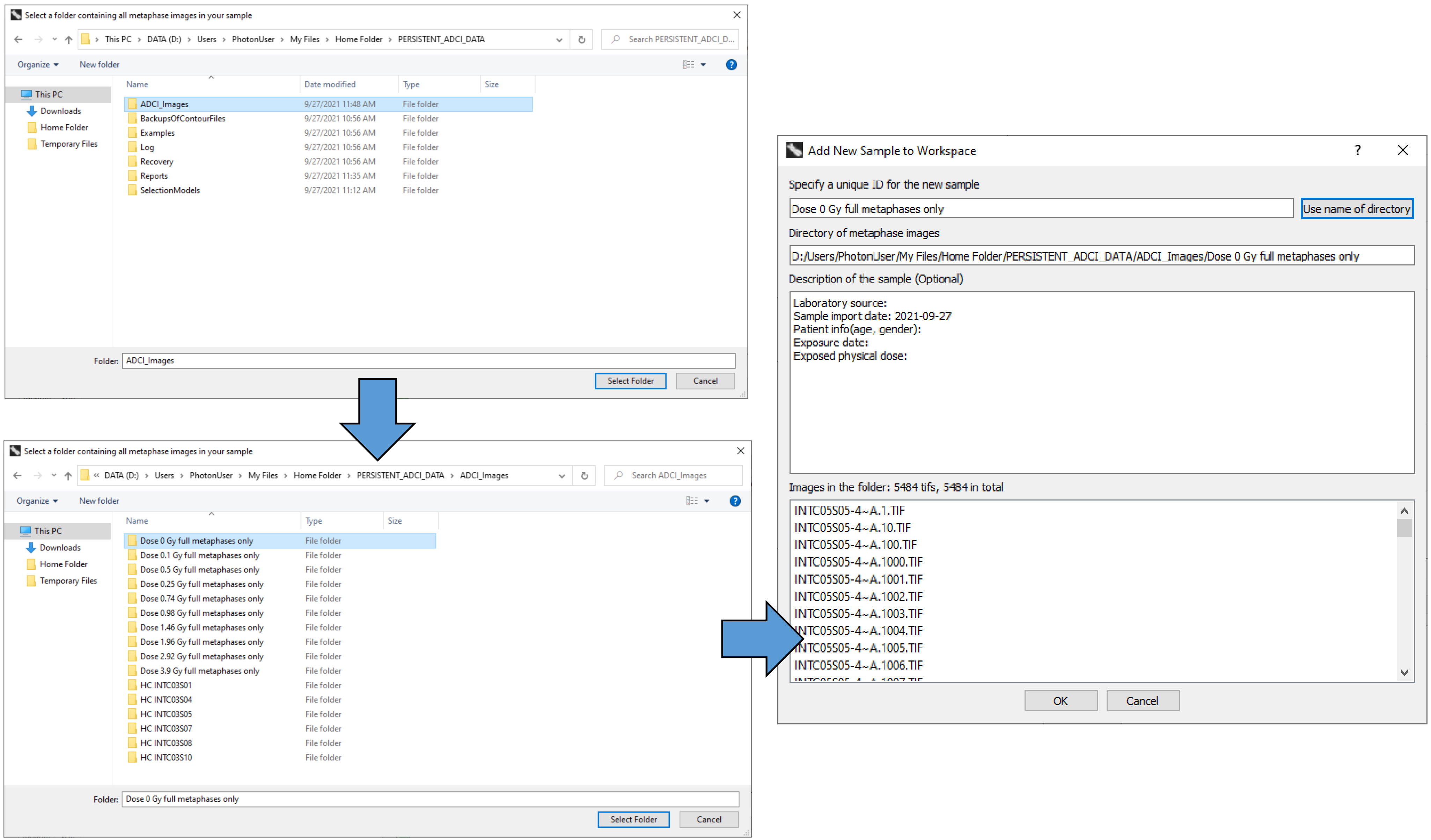

### Add Samples to the Process Queue

Time Required:
< 10 seconds

- After adding all desired samples to ADCI, samples must be processed (meaning ADCI examines metaphase images and locates dicentric chromosomes).
- Samples are added to the process queue, then ADCI is instructed to process all samples in the process queue.

- A minimal workflow for a typical ADCI session is:
  - Add samples
  - Process samples
  - Create a calibration curve
  - Perform dose estimation
  - Generate and review reports

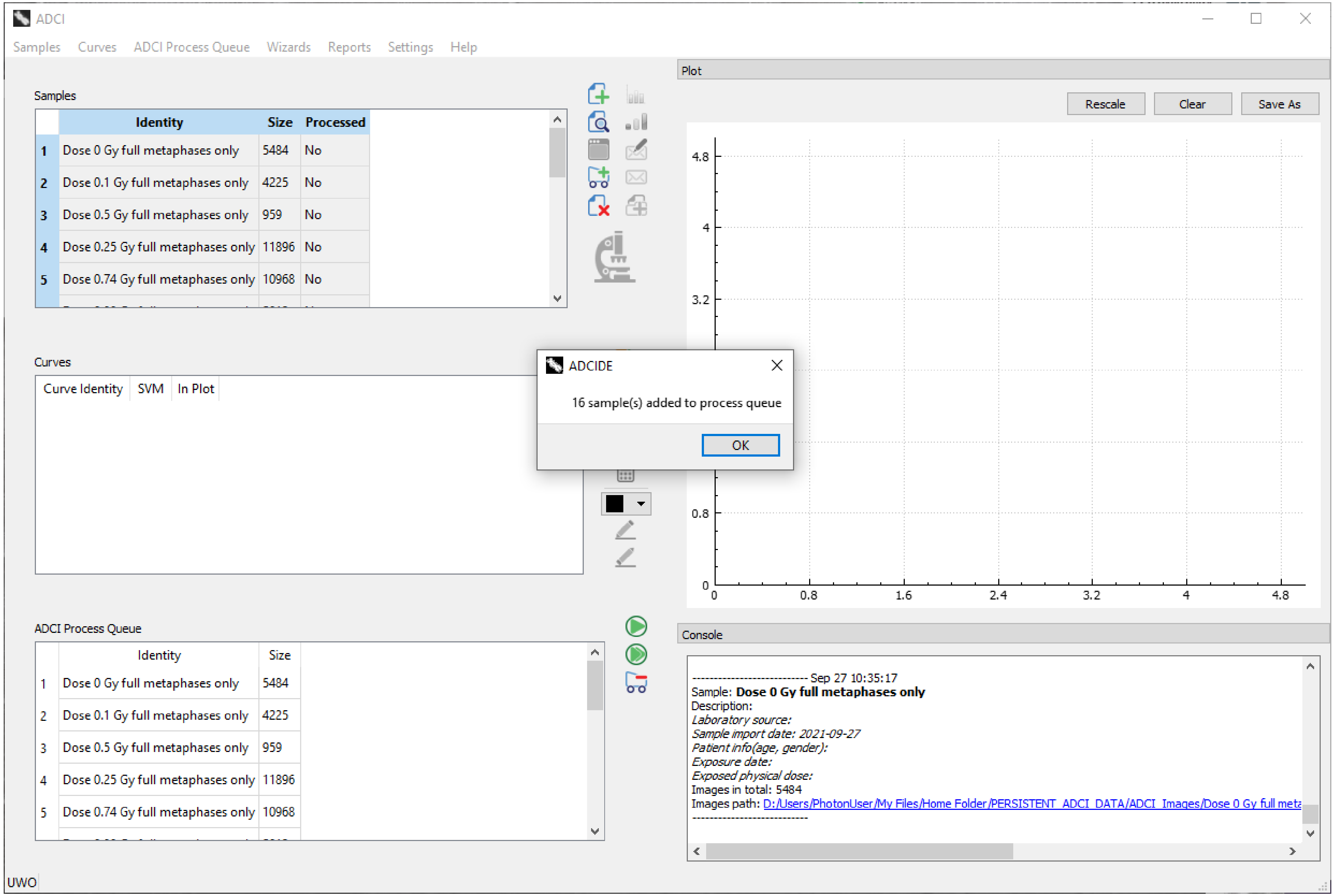

### Process Samples and Save Results

- After all samples in the process queue have been processed, results should be saved as “adcisample” files.
- Results are saved to cloud storage (“PERSISTENT_ADCI_DATA”) by default, allowing them to persist from one streaming session to the next.

- Sample processing is typically the most time-consuming step in an analysis, therefore selecting an appropriate (set of) streaming instance hardware configuration(s) on which to process samples is important when attempting to maximize efficiency.
- The approximate image processing rates displayed in the “Time Required” section can be applied to the number of metaphase images in a new analysis to obtain an estimate of the time required on different configurations.

<u>Time Required:</u>

**Image processing rates**:
standard[a]: 19.58 images/min
standard x5[b]: 97.53 images/min

**7 calibration samples (7500 images)**:
standard[a]: 6 hr, 23 min
standard x5[b]: 1 hr, 17 min

**10 test samples (8500 images)**:
standard[a]: 7 hr, 14 min
standard x5[b]: 1 hr, 27 min

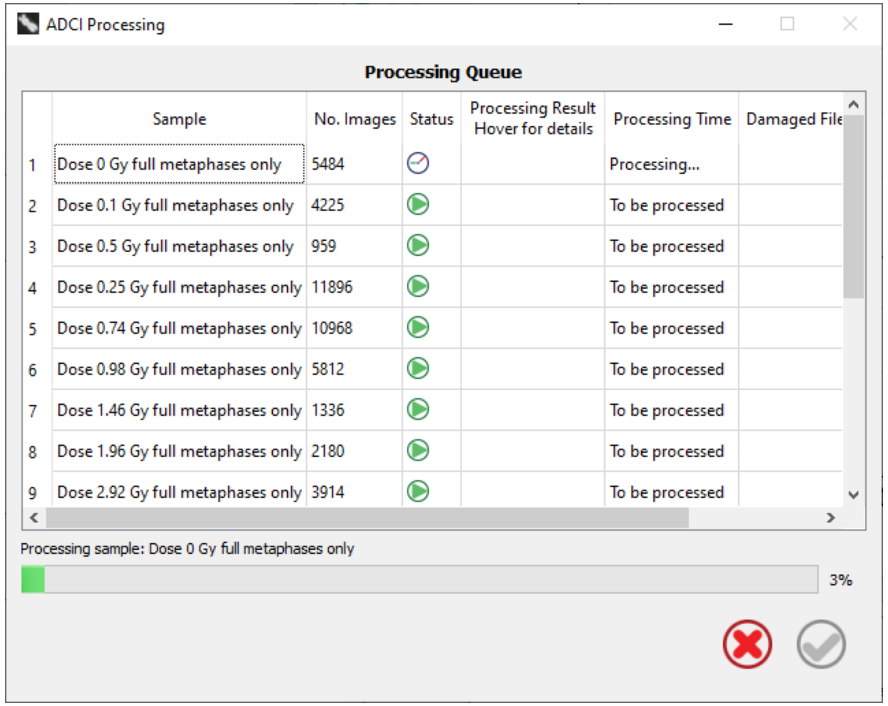


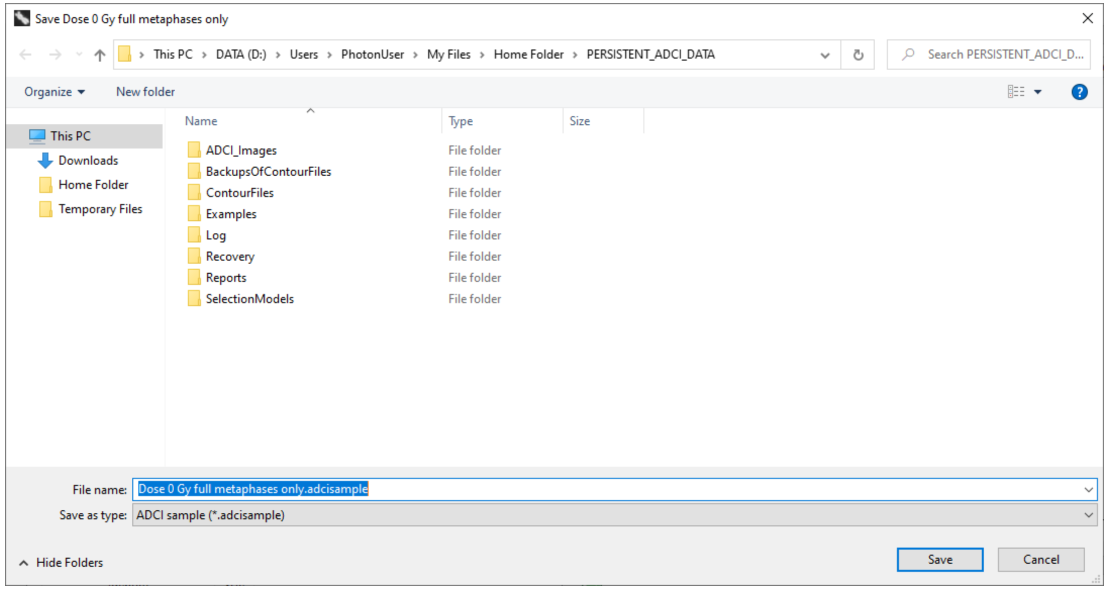


[a] Refers to a “stream.standard.medium” AppStream 2.0 hardware configuration (see manuscript, Table 1).
[b] Samples distributed among five “stream.standard.medium” instances executed in parallel.

### Optimal Image Selection Model Generation

<u>Time Required:</u>

**7 calibration samples (7500 images)**:

Leave-one-out evaluation method: 3 hr, 53 min

Other evaluation methods: 1 hr, 34 min

- Image selection models exclude metaphase images with suboptimal image quality or chromosome morphology from further analysis.
- Models are comprised of image exclusion filters, filter weights (optional), and an image ranking method (optional).
- A base set of image selection models is available, however new image selection models can also be created.
- The Image Selection Optimization Wizard generates a large set of models based on a user-specified search-space. Models generated in this way are applied to a set of calibration samples, and models are ranked according to one of three evaluation methods:
  - p-value of Poisson fit
  - curve fit residuals
  - leave-one-out cross validation

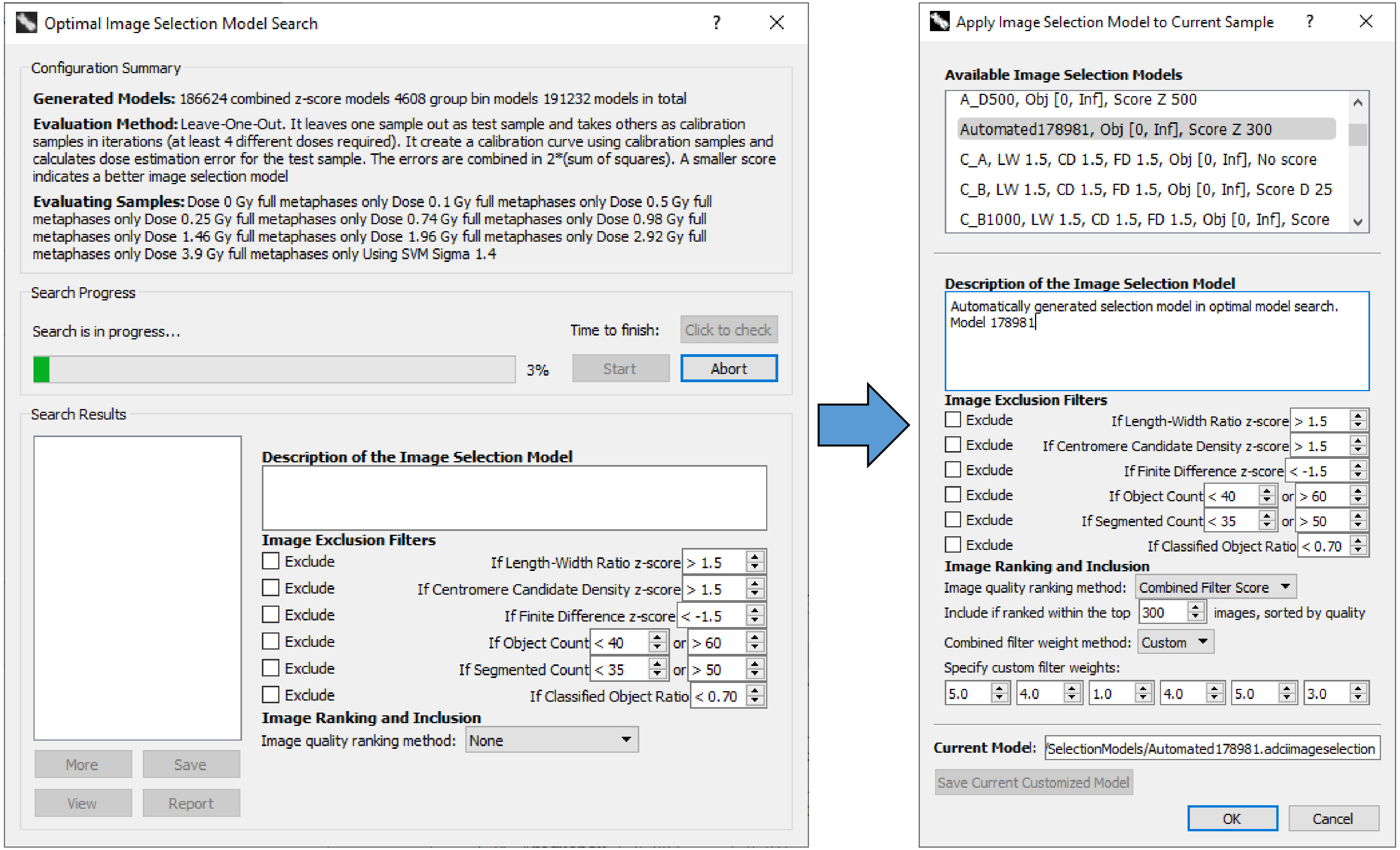

### Calibration Curve Generation

**Time Required:**
**5 - 30 min**

- The two dialogs (to the right) are prepopulated upon completion of the Calibration Curve and Dose Estimation wizards, respectively.
- While proceeding through the Calibration Curve wizard, an image selection model is specified and automatically applied to calibration samples (model “A_B” in this example). The chosen image selection model becomes associated with the calibration curve and is then automatically applied to test samples when dose estimation is performed.
- The machine learning tuning parameter “sigma” influences the number of DCs found in metaphase images (a higher sigma value results in a higher number of chromosomes flagged as DCs). It is been our experience that sigma values of 1.4 or 1.5 contribute to the most accurate dose estimation results.

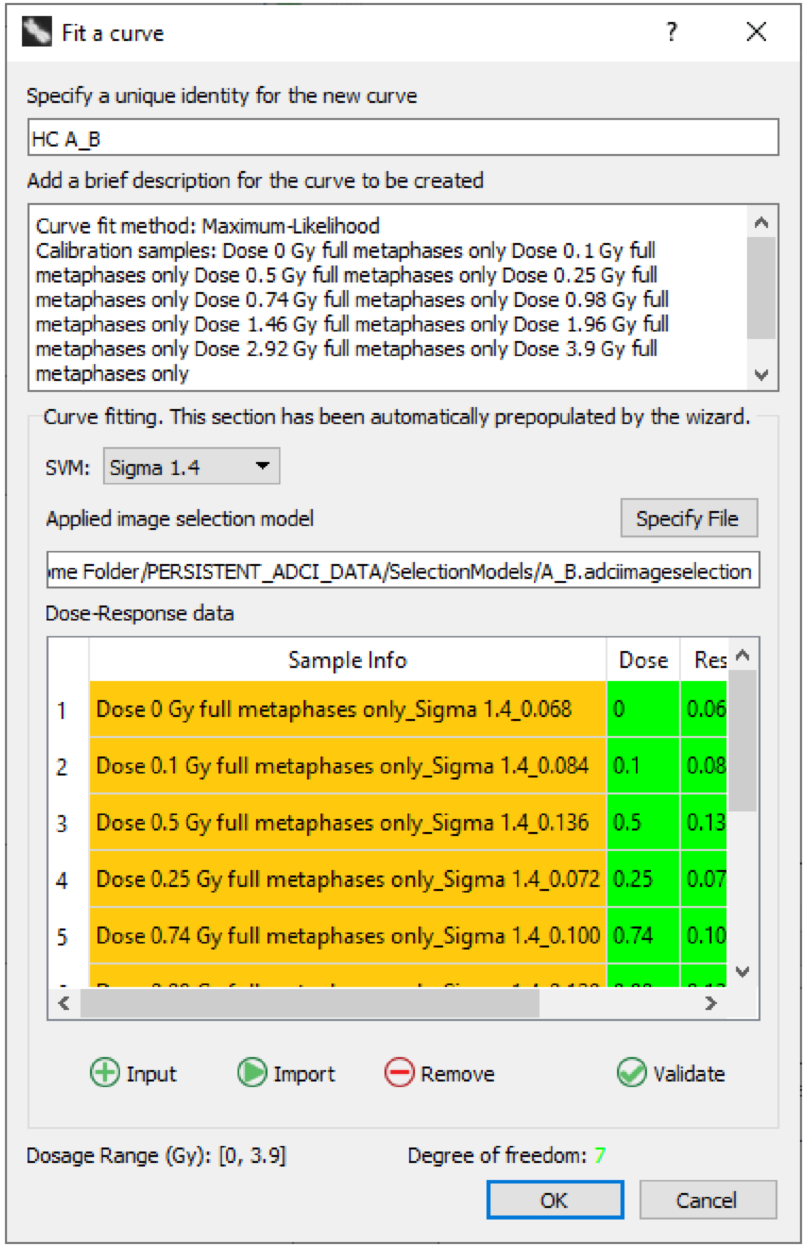


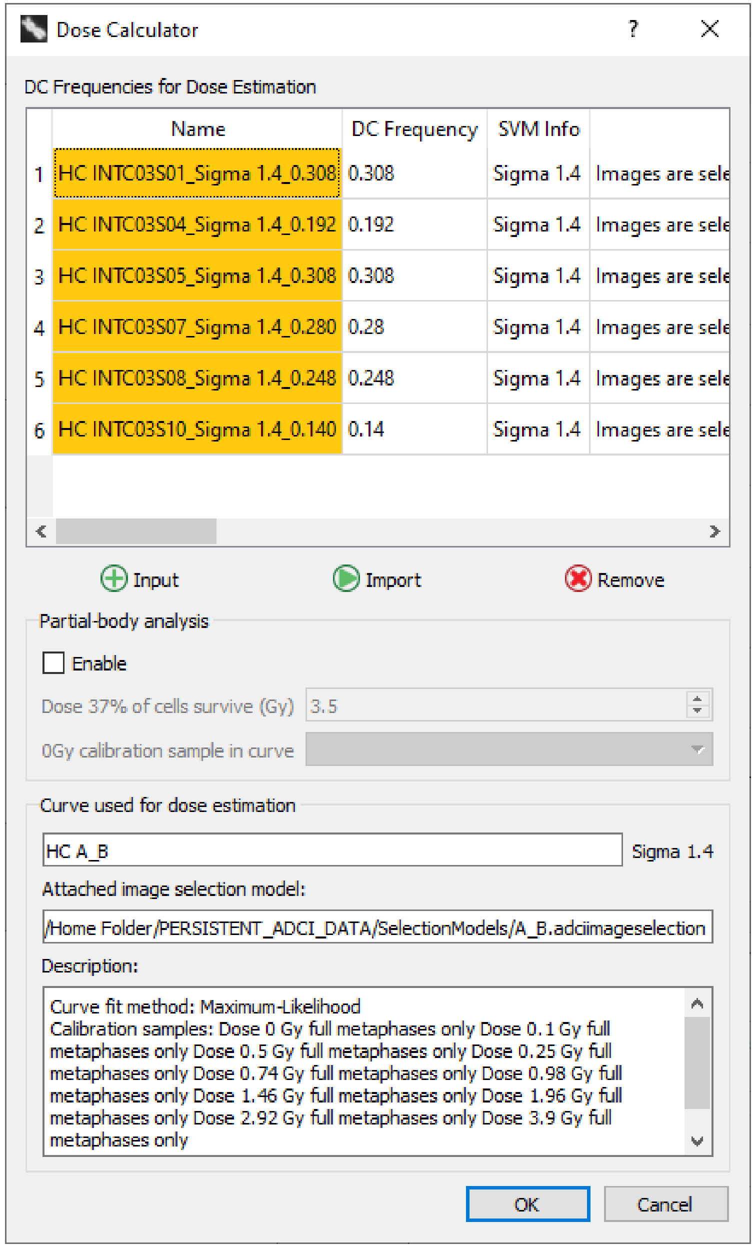

### Dose Estimation

**Time Required:**
5 - 30 min

- The same sigma value selected during calibration curve generation should be applied when filling out the dose estimation wizard.
- After completing the wizard and executing dose estimation, results appear in the ADCI console (bottom-right), and are plotted (top-right). Confidence intervals can optionally be displayed in the plot.
- DC frequencies of each calibration sample are plotted as black diamonds.
- Colored dashed lines represent the DC frequency and estimated dose of each test sample.
- The console output and plot - along with some additional information - can be saved as a Dose Estimation report (see next slide).

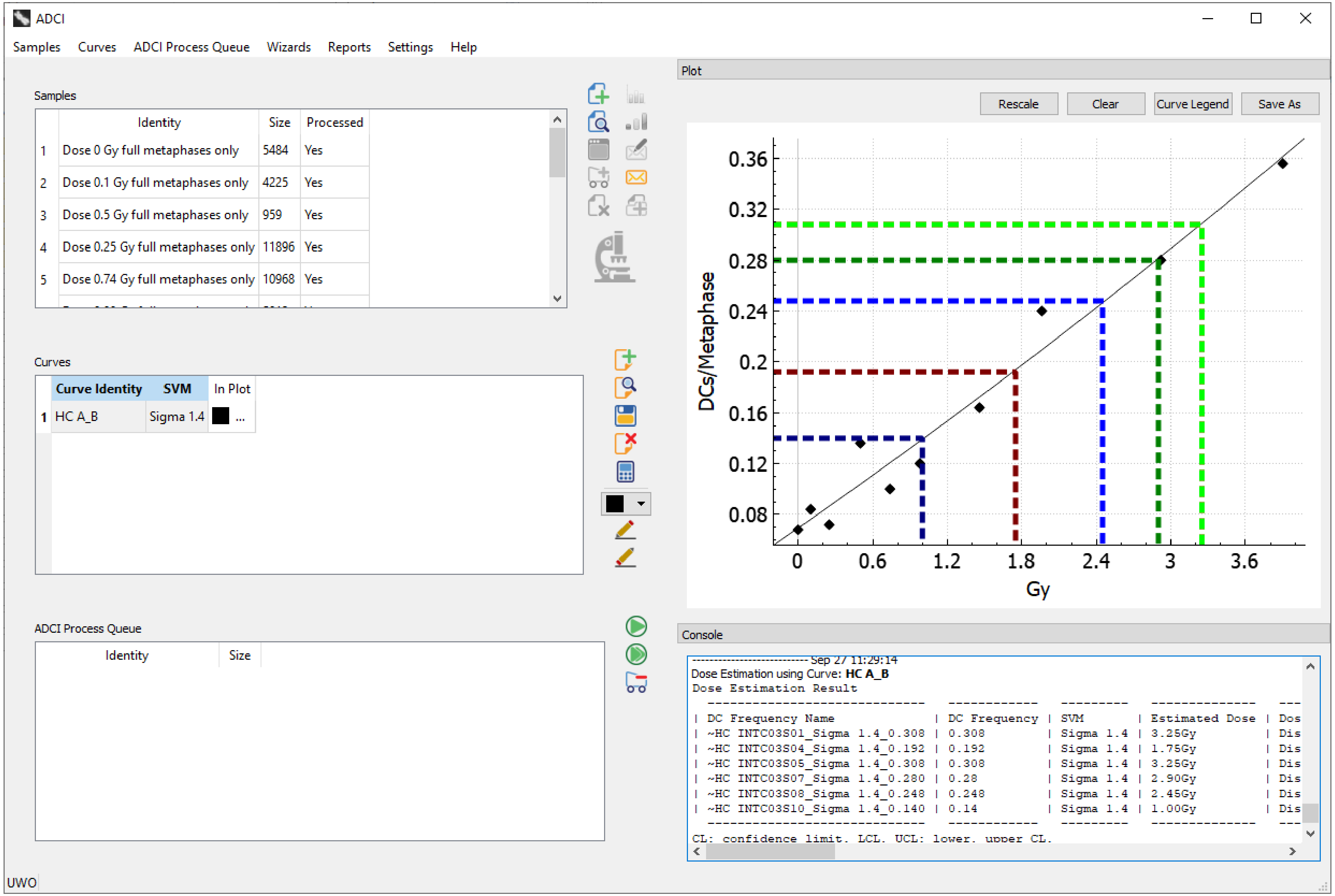

### Report Generation and Review

**Time Required:**
Report generation: < 10 seconds
Report review: 10 – 120 min

#### Dose Estimation Report

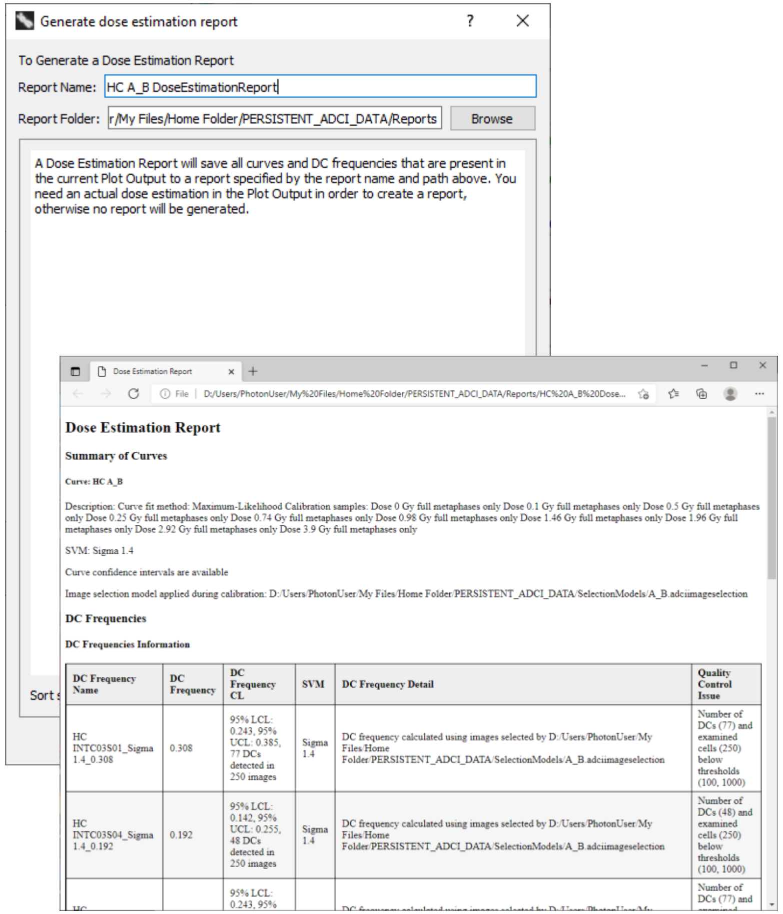


#### Calibration Curve Report

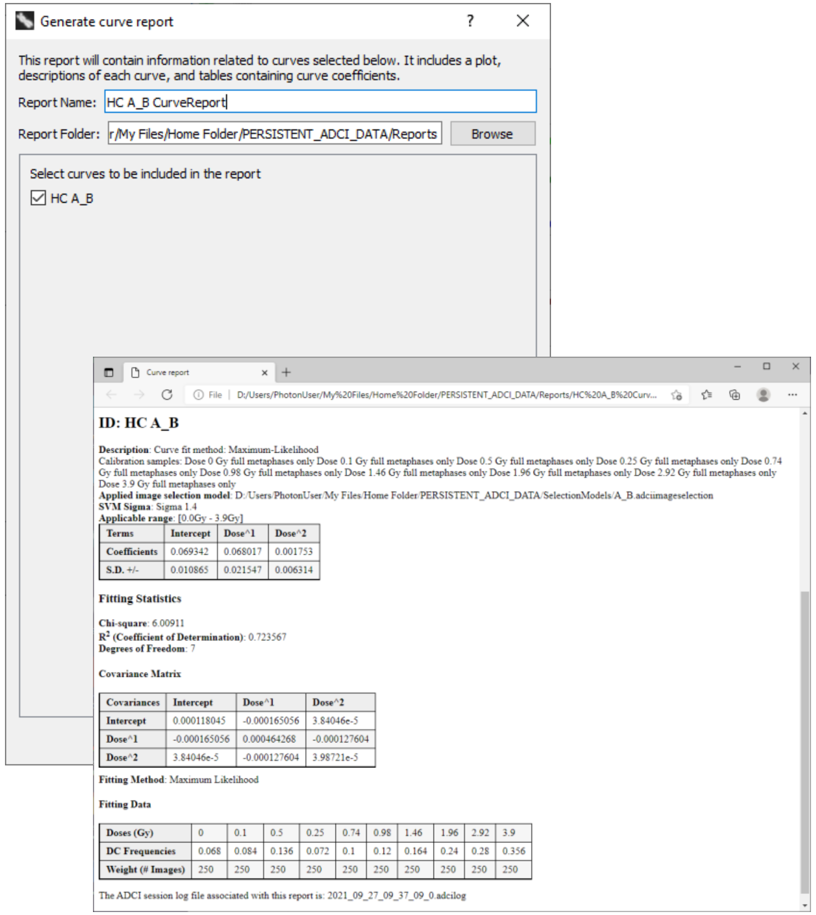


#### Sample Report

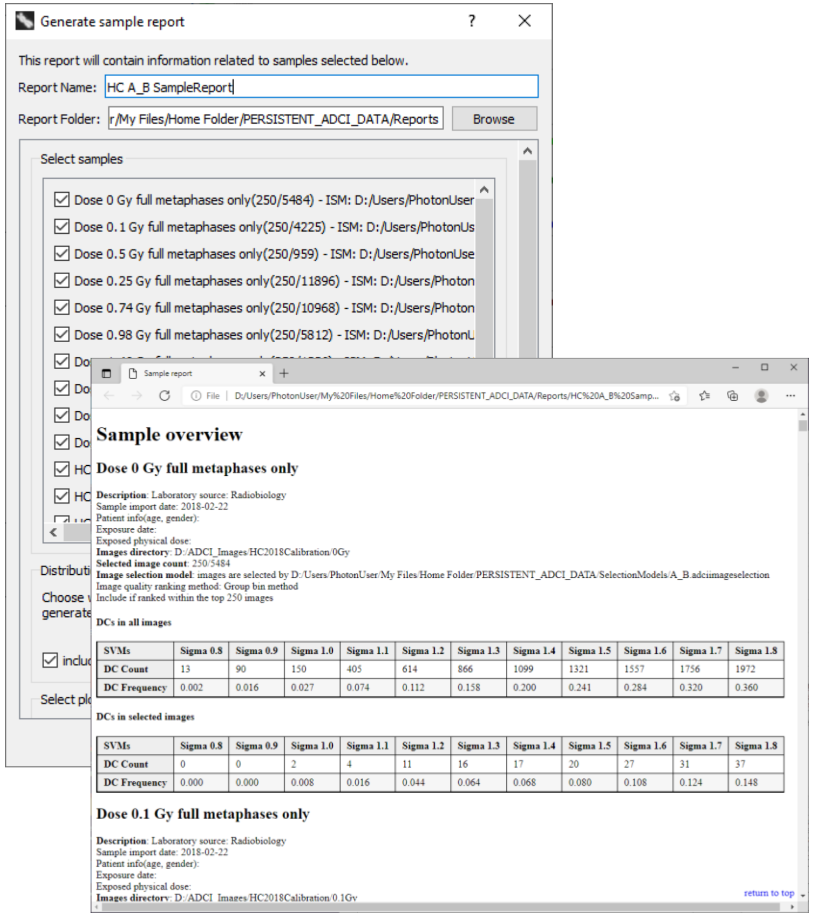


Reports are HTML pages (saved to disk and viewable offline) consisting of text, tables, and images that preserve data for future consultation. They may also be used to replicate results in the future as, for example, linear-quadratic terms and covariance matrix are present in the Calibration Curve Report, allowing the same curve to be manually input to ADCI if the saved curve file cannot be located. Reports are saved to cloud storage and can be downloaded using to a local system using the Javascript web app.

### Additional Information

ADCI online manual (wiki):

https://adciwiki.cytognomix.com

Introduction and access to demonstration version:

https://radiation.cytognomix.com

ADCI video protocol in the Journal of Visualized Experiments (JoVE):

https://doi.org/10.3791/56245

Dicentric chromosome classification by machine learning:

https://cytobiodose.cytognomix.com
